# Comprehensive single cell analyses of the nutritional environment of intracellular *Salmonella enterica*

**DOI:** 10.1101/2020.11.01.364018

**Authors:** Jennifer Röder, Pascal Felgner, Michael Hensel

**Affiliations:** Abteilung Mikrobiologie, Universität Osnabrück, Osnabrück, Germany; CellNanOs – Center of Cellular Nanoanalytics, Fachbereich Biologie/Chemie, Universität Osnabrück, Osnabrück, Germany

**Author notes:** Address for correspondence:* Michael Hensel, Abteilung Mikrobiologie, Fachbereich Biologie/Chemie, Universität Osnabrück, Barbarastr. 11, 49076 Osnabrück, Germany.

## Abstract

The facultative intracellular pathogen *Salmonella enterica* Typhimurium (STM) resides in a specific membrane-bound compartment termed the *Salmonella*-containing vacuole (SCV). STM is able to obtain all nutrients required for rapid proliferation, although being separated from direct access to host cell metabolites. The formation of specific tubular membrane compartment, called *Salmonella*-induced filaments (SIFs) are known to provides bacterial nutrition by giving STM access to endocytosed material and enabling proliferation. Additionally, STM expresses a range of nutrient uptake system for growth in nutrient limited environments to overcome the nutrition depletion inside the host. By utilizing dual fluorescence reporters, we shed light on the nutritional environment of intracellular STM in various host cells and distinct intracellular niches. We showed that STM uses nutrients of the host cell and adapts uniquely to the different nutrient conditions. In addition, we provide further evidence for improved nutrient supply by SIF formation or presence in the cytosol of epithelial cells, and the correlation of nutrient supply to bacterial proliferation.

## Introduction

*Salmonella enterica* is an invasive, facultative intracellular pathogen causing diseases ranging from gastroenteritis to systemic typhoid fever. With 93.8 million foodborne diseases and 155,000 deaths per year, *Salmonella* is a major public health problem (Majowicz et al., 2010). In some cases, *Salmonella* infections are life-threatening and require appropriate and effective antibiotic therapy. The development of multidrug-resistant *Salmonella* serotypes has a major impact on the effectiveness of antibiotic therapy, therefore new anti-infective strategies are required (Eng et al., 2015). The understanding of the metabolism of *Salmonella*, which is prerequisite for bacterial viability and virulence, could be new targets for antimicrobial therapies.

*Salmonella enterica* serovar Typhimurium (STM) resides in a specialized membrane-bound compartment termed *Salmonella*-containing vacuole (SCV), which allows survival and proliferation of the bacteria within host cells (Haraga et al., 2008). Essential for intravacuolar survival is the function of the type III secretion system (T3SS) encoded by genes on *Salmonella* pathogenicity island 2 (SPI2) (Hensel et al., 1998). The SPI2-T3SS translocates a set of effector proteins across the SCV membrane into the host cytosol in order to manipulate host cell functions. Mutant strains defective in SPI2-T3SS are attenuated in systemic virulence and show reduced intracellular replication (Hensel et al., 1995; Hensel et al., 1998). In addition, translocation of SPI2 effector proteins induces the formation of specific tubular membrane compartments growing out of the SCV, called *Salmonella*-induced filaments (SIFs) (Brumell et al., 2001b; Drecktrah et al., 2007; Rajashekar et al., 2008; Figueira and Holden, 2012). Most important SPI2-T3SS effector is SifA, a lack of this effector leads to defects in remodeling of the host cell endosomal system and maintenance of intact SCV (Stein et al., 1996; Beuzon et al., 2000). A Δ*sifA* strain is attenuated in replication in macrophages (Beuzon et al., 2000; Brumell et al., 2001a) and results to hyper-replication of STM in the nutrient-rich cytosol of epithelial cells (Knodler, 2015). Replication indicated in the cytosol suggests an alternative niche for replication, in addition to the SCV, and a bimodal lifestyle has been suggested (Malik-Kale et al., 2012). Inside host cells, STM has to resist various antimicrobial strategies of the host, such as antimicrobial peptides, reactive oxygen species, acidification of vacuoles, and nutrient restriction (Flannagan et al., 2009). The limitation of nutrients by the innate immune response of the host, referred as nutritional immunity, is a challenge for intracellular pathogens. Concentrations of essential elements such are iron and zinc decrease rapidly during infection to starve invading pathogens (Hennigar and McClung, 2016). Conversely, pathogens have acquired various mechanisms to bypass the innate nutritional immunity of the host, e.g. by developing a number of transporters and binding proteins (Hennigar and McClung, 2016). By deploying such functions, STM is able to subvert these defense systems and successfully colonize the host cell (Flannagan et al., 2009).

STM resides within the nutrient-poor SCV, but is able to obtain all nutrients required for rapid proliferation, although being separated from direct access to host cell metabolites. The induced formation of an extensive SIF network by STM is described to support bacterial nutrition and enables proliferation by giving STM access to endocytosed material (Liss et al., 2017). In addition, there are a range of nutrient uptake system for growth in nutrient-poor environments to overcome the nutrition depletion. The SCV is characterized by a mildly acidic pH, low magnesium (Mg^2+^), low iron (Fe^2+^), and low phosphate (Pi) content, and the presence of antimicrobial peptides (Deiwick and Hensel, 1999; Lober et al., 2006). Several analyses of entire intracellular STM population by transcriptomics or proteomics have shown up-regulation of various nutrient uptake systems within the intracellular environment, allowing STM to proliferate in the nutrient-limited environment (examples in Kroger et al., 2013; Liu et al., 2015; Noster et al., 2019). However, relatively little is known regarding the nutritional and metabolic requirements of *Salmonella* during infection. Intracellular STM behave highly heterogenous and form subpopulations with rather diverse physiological properties (reviewed in Helaine and Holden, 2013; Bumann, 2015). In addition, the respective niches vary in different host cells and require optimal, unique nutritional and metabolic adaptation of STM. These metabolic flexibilities allow STM to colonize various environmental niches and host cells.

Here, we investigated the nutritional environment of intracellular STM in various host cells at a single cell level by using dual fluorescent reporters in combination with flow cytometry (FC). We determined the contribution of SPI2-T3SS activity, the nutritional effects of cytosolic lifestyle, and influence of bacterial proliferation to study heterogeneity of intracellular STM in relation to nutrient supply and acquisition. Since in some cases, e.g. in immunocompromised patients, STM can cause invasive non-typhoidal Salmonellosis, we also addressed the rather unknown environment in primary human macrophages. We investigated the nutritional conditions for STM in primary human macrophages. The improved understanding of the nutritional environments in mammalian host cells, and adaptation of STM metabolism could contribute to the development of new targets for antimicrobial therapies.

## Results

### Single cell-based approach to determine nutrient availability of intercellular STM

Here we established reporter systems to precisely characterize the nutritional environment of intracellular STM at single cell level. For this, we generated dual fluorescence reporter plasmids with constitutive expression of DsRed, and sfGFP under control of the promoter of genes for acquisition of specific nutrients. The overall procedure is described in **Figure 1**. STM strains containing reporter plasmids were cultured in defined media with varying amounts of the nutrients to be analyzed. Fluorescence intensities of STM were measured by flow cytometry (FC) to determine the sfGFP intensity as function of various nutrient concentrations. Subsequently, epithelial cells and macrophages were infected with STM containing the dual fluorescence reporters, lysed and intracellular released bacteria measured by FC. DsRed fluorescence allowed detection of STM by FC in complex samples such as lysates of host cells, and dilution of slowly maturing DsRed by cell division served as proxy for the proliferation rate of STM. A correlation of the sfGFP intensities was performed, which provided values for nutrient concentration available. Since this was done at single cell level, population changes over time can be observed and correlated with the intracellular habitat, e.g., vacuolar vs. cytosolic presence in HeLa cells. In analyses to distinguish responses of SCV-bound from those of cytosolic STM, P_EM7_::DsRed was replaced by P_*uhpT*_::DsRed as sensor for glucose-6-phosphate (G6P) present in the host cell cytosol, as previously reported (Röder and Hensel, 2020a).

**Figure 1:**
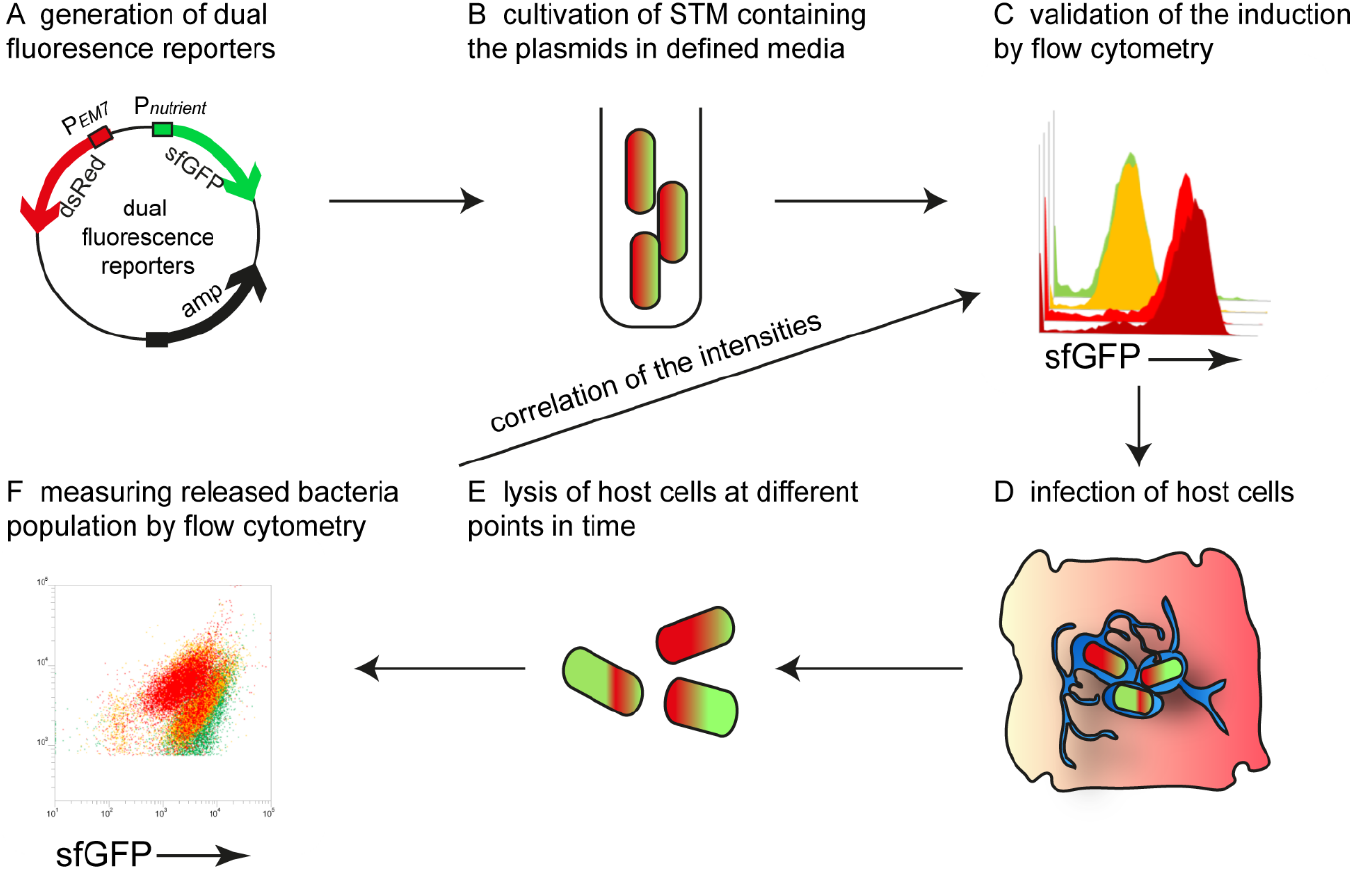
Intracellular STM as sensor to determine nutrient availability in host cells. A) Dual fluorescence reporter plasmids feature constitutive expression of DsRed, and expression of sfGFP under control of promoters of genes involved in nutrient uptake or metabolism. B) For *in vitro* validation of reporters, STM was cultured in defined media with various concentration of the nutritional factor of interest. C) Gating of DsRed-positive cells and quantification of sfGFP intensity by flow cytometry (FC). D) STM strains harboring confirmed reporter plasmids were used to infect host cells. E) At various timepoints p.i., the infected host cells were lysed. F) The released bacteria were analyzed by FC. A comparison of the sfGFP intensities was performed, providing the nutrient concentration available for STM at single cell level.

Reporter expression by plasmid-borne fusions may be affected by the copy number of the vector. With the example of the iron-responsive P_*sitA*_::sfGFP fusion, we analyzed the effect of single, low, or mid copy number of the vector(**Figure S 1**). Under iron-limiting conditions, the X-mean sfGFP intensities of STM increased with copy number of the vector. However, the distribution of the STM population was comparable. As further test for specificity of the reporter system, in analysed induction of representative fusions P_*bioA*_::sfGFP or P_*corA*_::sfGFP in corresponding Δ*bioA* or Δ*corA* background, respectively, in comparison the WT background (**Figure S 2**). The sfGFP intensities in mutant background were increased for both fusions, indicating that the reporters faithfully sense the intracellular conditions in STM and starvation for nutrients.

### Potassium is not limited in host cells

Potassium ions (K^+^) are essential for the physiology of bacteria and involved in almost all aspects of growth. Potassium is the most abundant cation and its homeostasis in STM is regulated by three major K^+^ transporters, i.e. low affinity transporters Kup and Trk, and high affinity uptake system Kdp (Su et al., 2009). We generated a dual fluorescence reporter using the *kdpA* promoter to analyze potassium availability of STM in host cells. Due to the described induction of the *kdp* system in environments with low K^+^ (5 mM or less) *(Su et al., 2009)*, we tested *in vitro* induction by using an MKM medium and added various concentration of KCl. In media with 10 mM KCI, no induction of the reporter was detected, but concentrations lower than 1 mM KCI led to strong induction of P_*kdpA*_::sfGFP (**Figure 2**A). In HeLa cells infected with STM WT, we did not detect induction of the potassium reporter, indicating that at least concentrations between 5-10 mM K^+^ are available in intracellular habitats of HeLa cells (**Figure 2**B). Only at 16 h p.i., a small induced population of ca. 7% of STM WT in RAW264.7 macrophages indicated limitation (**Figure 2**A, black frame). This induction corresponded to concentrations of less than 1 mM K^+^ under *in vitro* conditions.

In line with FC data, micrographs of infected HeLa cells did not reveal sfGFP signals (**Figure 2**D), while a rather heterogeneous distribution of a low number of P_*kdpA*_-positive STM was observed in RAW264.7 macrophages (**Figure 2**E). Therefore, K^*+*^ is not limited for STM in host cells.

**Figure 2:**
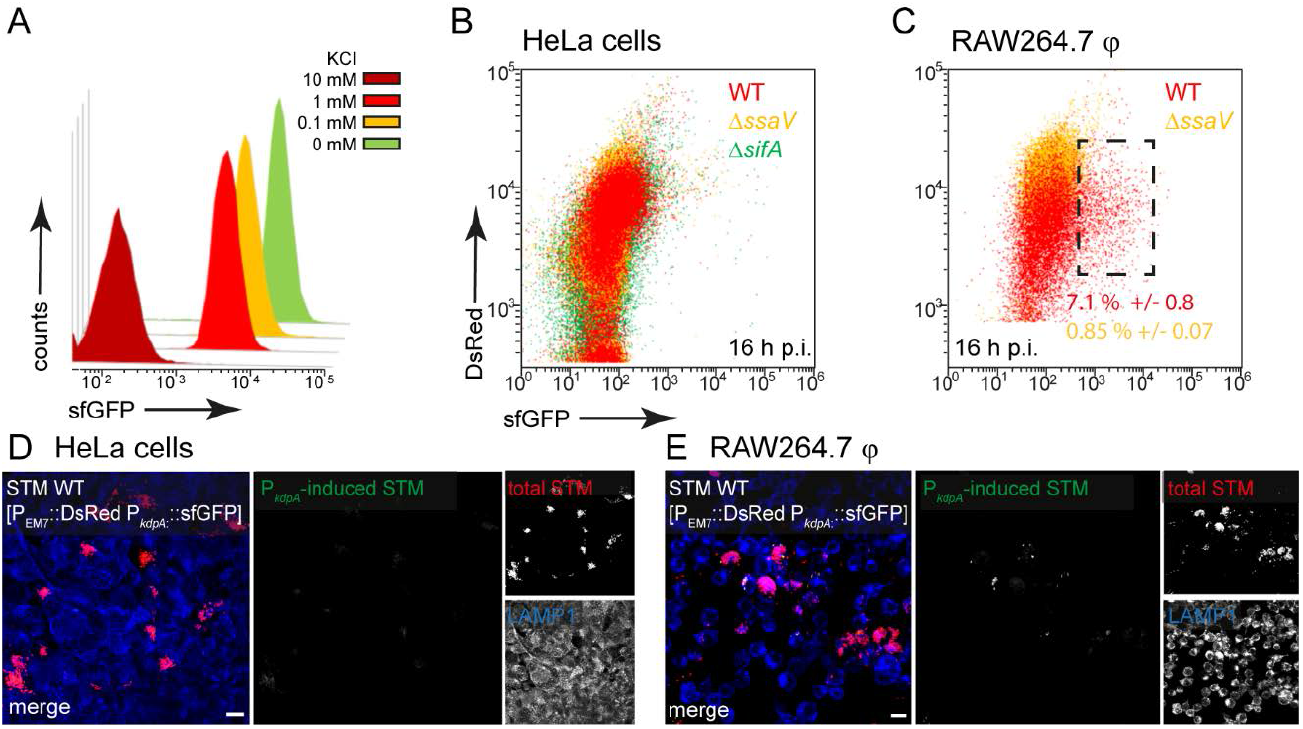
A dual fluorescence reporter for measuring the intracellular potassium availability for STM in HeLa cells or RAW264.7 macrophages. STM harboring p5080 for constitutive expression of DsRed, and sfGFP under control of P_*kdpA*_ was cultured in MKM minimal medium with various amounts of KCI. STM WT [p5080] was grown overnight (o/n) in PCN containing 25 mM Pi (PCN (25)), pH 7.4, diluted 1:31 in MKM with various concentrations of KCI as indicated and subcultured for 3.5 h. Samples were collected after 3.5 h, sfGFP intensity of P_*kdpA*_-induced bacteria was determined by FC (A). sfGFP intensities of P_*kdpA*_-positive bacteria of a representative experiment are shown. The values for induction of the potassium reporter were derived from at least three independent experiments. Host cells were infected at MOI 5 with STM WT (red), Δ*ssaV* (orange), or Δ*sifA* (green) strains as indicated, each containing the potassium reporter p5080. HeLa cells (B), or RAW264.7 macrophages (C) were lysed 16 h p.i., released STM were fixed, and subjected to FC to quantify sfGFP intensities of P_*kdpA*_-positive STM. P_*kdpA*_-positive bacterial populations of a representative experiment from three biological replicates are shown. HeLa cells (D) and RAW264.7 macrophages (E) were infected with STM WT [p5080] and immuno-stained against LAMP1 (blue) for labelling of late endosomal and lysosomal membranes. Cells were imaged 16 h p.i., and overview images show heterogeneous sfGFP (green) intensities of STM. Representative infected cells indicate DsRed (red) and sfGFP (green) fluorescence signal for STM. Scale bars, 10 μm.

### The access of STM to magnesium and zinc is heterogeneous

STM possesses three transport systems for Mg^2+^, i.e. CorA, MgtA, and MgtB. The *corA* gene is described as constitutively expressed but was up-regulated in media with low Mg^*2+*^ concentrations that mimic conditions within the SCV, and CorA mediates both uptake and release of Mg^*2+*^ (Hmiel et al., 1989; Kroger et al., 2013). Mg^2+^ is involved in several stages of STM infection and virulence, for example, low extracellular Mg^2+^ levels were shown to activate SPI2 gene expression (Deiwick and Hensel, 1999; Papp-Wallace et al., 2008). We generated a magnesium reporter with the promoter of *corA*, which showed a continuous increase in sfGFP intensity with decreasing Mg^2+^ concentration in PCN medium. Concentrations of MgSO_4_ lower than 10 μM led to sfGFP intensity above 800 RFI (**Figure 3**AB).

**Figure 3:**
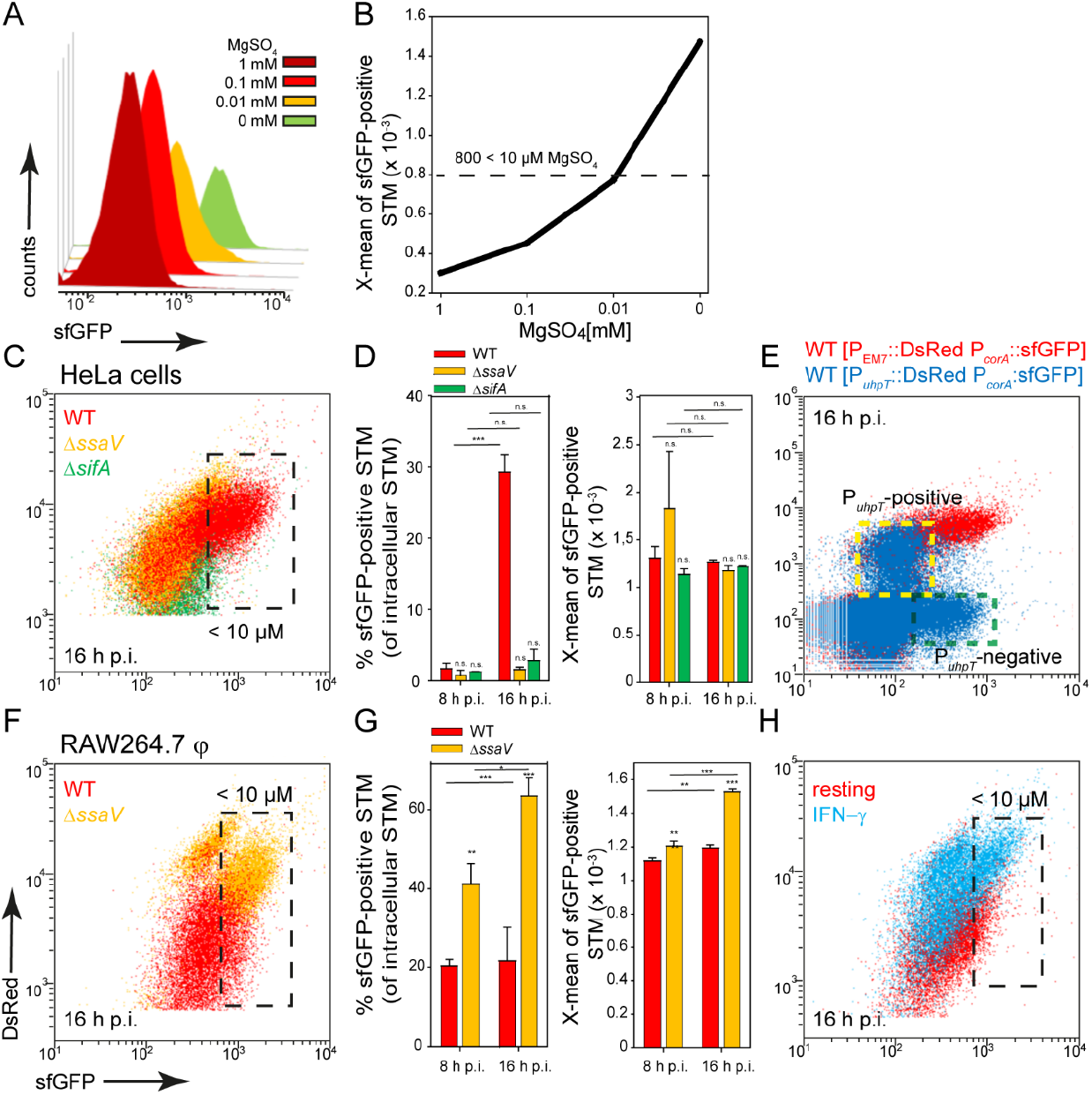
A dual fluorescence reporter for measuring the intracellular magnesium availability for STM in HeLa cells or RAW264.7 macrophages. STM harboring p5078 for constitutive expression of DsRed, and sfGFP under control of P_*corA*_ was cultured in in PCN minimal medium with various amounts of MgSO_4_. STM WT [p5078] was grown o/n in PCN (25), pH 7.4, diluted 1:31 in fresh PCN (1), pH 7.4 with various concentrations of MgSO_4_ as indicated and subcultured for 3.5 h. Samples were collected after 3.5 h (A) and X-means of sfGFP intensity of P_*corA*_-induced bacteria was determined by FC (B). sfGFP intensities above 800 RFI indicate MgSO_4_ concentrations lower than 10 μM. sfGFP intensities of P_*corA*_-positive bacteria of a representative experiment are shown. The values for the induction of the magnesium reporter were derived from at least three independent experiments. Host cells were infected with STM WT (red), Δ*ssaV* (orange) and Δ*sifA* (green) strains as indicated, each containing the magnesium reporter p5078 at MOI of 5. HeLa cells (C, D) or RAW264.7 macrophages (F, G) were lysed at 8 h or 16 h p.i., released STM were fixed, and subjected to FC to quantify sfGFP intensities of P_*corA*_-positive STM. Representative quantification of population size and X-mean sfGFP intensities of P_*corA*_-positive population for STM WT, Δ*ssaV* and Δ*sifA* in HeLa cells (D), or in RAW264.7 macrophages (G) at 8 h and 16 h p.i. Mean values and standard deviations of P_*corA*_-positive bacterial populations from triplicates of a representative experiment from three biological replicates are shown. Statistical analyses were performed by one-way ANOVA for STM WT compared to mutant strains, or between time points, and are expressed as: n.s., not significant; *, p <0.05; **, p < 0.01; ***, p < 0.001. E) HeLa cells were infected at MOI 5 with STM WT harboring p5189 with DsRed under control of P_*uhpT*_ and P_*corA*_::sfGFP (blue). For comparison and gating of populations, HeLa cells were infected with STM WT harboring p5078 for constitutive expression of DsRed and P_*corA*_::sfGFP (red). Host cells were lysed 16 h p.i., released STM were fixed, and subjected to FC for quantification of P_*corA*_-positive bacteria and induction of P_*uhpT*_. Data for STM WT [p5078] and WT [p5489] (E), of a representative experiment from three biological replicates are shown. H) RAW264.7 macrophages were cultured in medium without (red) or with 5 ng x ml^-1^ IFN-γ (blue) for 24 h, and subsequently infected with STM WT [p5078] at MOI 5. Host cells were lysed at 16 h p.i., released STM were fixed, and subjected to FC to quantify P_*corA*_-positive STM. Data for STM WT [p5078] in resting RAW264.7 (red), or activated RAW264.7 (blue) of a representative experiment from three biological replicates are shown.

In infection experiments, about 95% of STM isolated from HeLa cells at 8 h p.i. showed a lower sfGFP intensity and therefore Mg^2+^ concentration above 10 μM available in the environment. Surprisingly, at 16 h p.i. only a small population of ca. 30% of STM WT showed a stronger induction comparable to the sfGFP intensity reached during growth in medium with 10 μM MgSO_4_, but this population was not detected in HeLa cell infected with STM Δ*ssaV* or Δ*sifA* (**Figure 3**C, black frame, D). Application of reporter P_*corA*_::sfGFP in combination with P_*uhpT*_::DsRed for cytosolic presence (Röder and Hensel, 2020a) clarified that induction of STM WT only occurred in the P_*uhpT*_-negative (vacuolar), but not in the P_*uhpT*_-positive (cytosolic) population (**Figure 3**E, indicated by yellow frame). Therefore, in the late SCV harboring STM WT, less than 10 μM Mg^2+^ is present, likely as consequence of the strong replication of STM WT in contrast to STM Δ*ssaV*. However, in the cytosol of the host cells no, or only very low induction was measured, indicating that more than 10 μM Mg^2+^ are available. In bacteria isolated from RAW264.7 macrophages, a slight induction was detected for almost all bacteria, but only ca. 20% of STM WT exhibited strong sfGFP intensities. In contrast, induction of 40% and 60% at 8 h p.i. and 16 h p.i., respectively, was recorded for intracellular STM Δ*ssaV* (**Figure 3**F, black frame, G). The isolation of bacteria from γ-interferon (INF-γ)-activated RAW264.7 macrophages resulted in mixed populations for both STM WT and Δ*ssaV* (**Figure 3**H, black frame), probably because of lower levels of replication and survival of STM (Rosenberger and Finlay, 2002). This also explained the higher DsRed intensity of single STM cells. The lower proliferation of STM Δ*ssaV* or STM WT isolated from IFN-γ-activated RAW264.7 macrophages resulted in accumulation of DsRed. In HeLa cells the difference is much smaller and indicated augmented proliferation of STM Δ*ssaV* in comparison to Δ*ssaV* in RAW264.7 macrophages. Based on these results, the SCV in RAW264.7 macrophages can be defined as magnesium-poor environment, but with ion concentrations mostly above 10 μM Mg^2+^. As the reporter showed a stronger induction in the STM Δ*ssaV* or in activated macrophages, it can be assumed that STM WT has more access to Mg^2+^ ions.

ZnuA is the periplasmic component of the high-affinity Zn^2+^ transporter ZnuABC, and *znuABC* expression is strongly induced in intracellular STM *(Ammendola et al., 2007)*. For the zinc reporter we used the promoter of *znuA* and investigated the induction in PCN medium with various concentrations of ZnSO_4_. The reporter showed a particularly strong induction in medium without ZnSO_4_, and relatively uniform sfGFP intensities between 1 μM and 1 mM ZnSO_4_. Without ZnSO_4_, X-means of sfGFP intensity above 600 RFI are measured (**Figure 4**AB). A comparable intensity was observed for about 10% and 50% of STM WT released from HeLa cells at 8 h or 16 h p.i., respectively. The STM WT showed the largest induced population compared to STM Δ*ssaV* and Δ*sifA* at 16 h p.i. (**Figure 4**C, black frame, D). When comparing the results with the double inducible reporter, again a stronger induction in P_*uhpT*_-negative (vacuolar) bacteria was observed. P_*uhpT*_-positive (cytosolic) bacteria showed only a slight induction of the reporter (**Figure 4**E, indicated by yellow frame). According to this, less than 1 μM Zn^2+^ is present in the SCV, while a higher Zn^2+^ concentration can be deduced as present in cytosol of HeLa cells.

**Figure 4:**
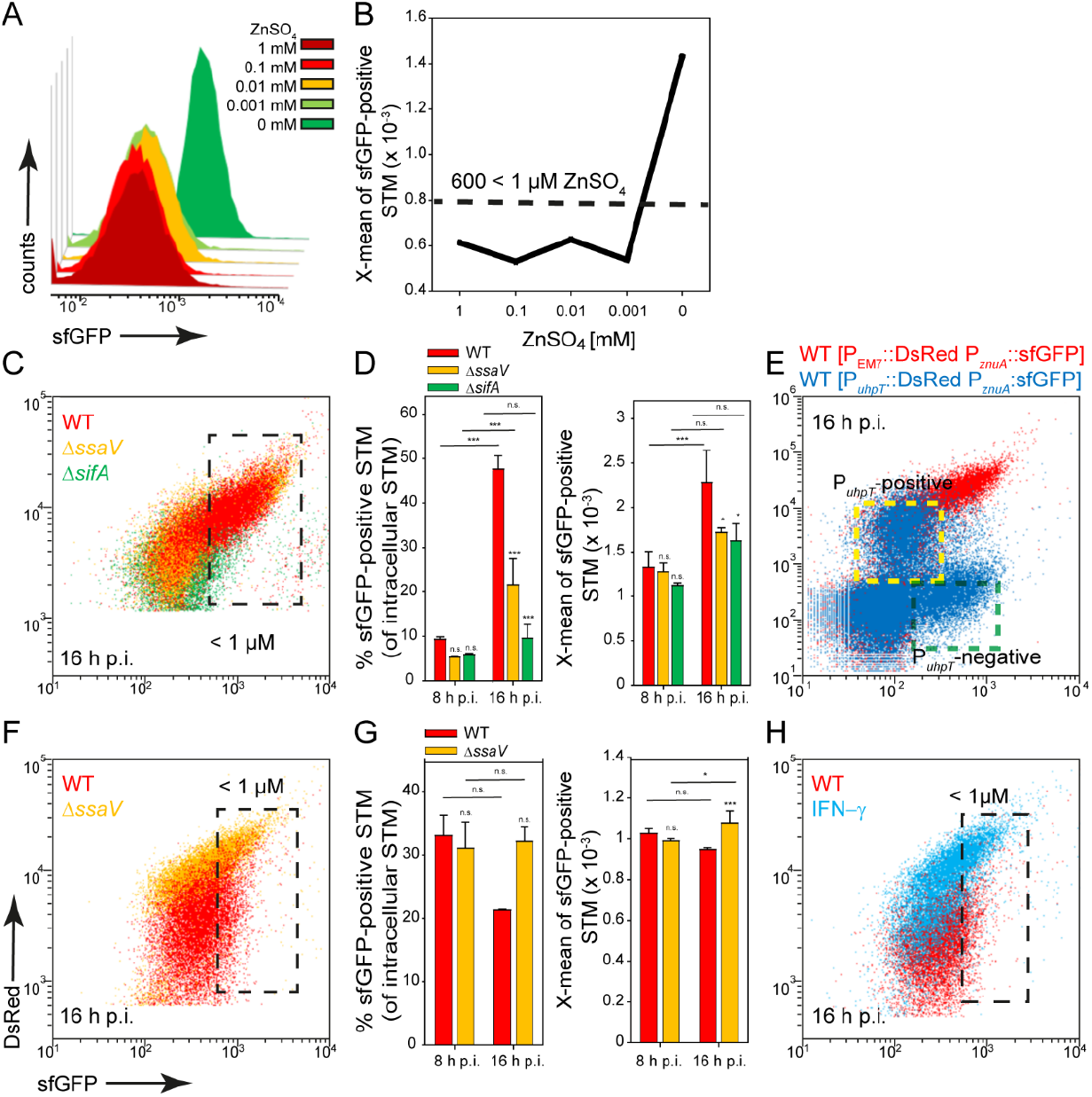
A dual fluorescence reporter for measuring the intracellular zinc availability for STM in HeLa cells or RAW264.7 macrophages. STM harboring p5003 for constitutive expression of DsRed, and sfGFP under control of P_*znuA*_ was cultured in in PCN minimal medium with various amounts of ZnS04. STM WT [p5003] was grown o/n in PCN (25), pH 7.4, diluted 1:31 in PCN (1), pH 7.4 with various concentrations of ZnSO_4_ as indicated, and subcultured for 3.5 h. Samples were collected after 3.5 h (A) and X-means of sfGFP intensity of P_*znuA*_-induced bacteria were determined by FC (B). sfGFP intensities above 600 RFI indicate ZnSO_4_ concentrations lower than 1 μM. sfGFP intensities of P_*znuA*_-positive bacteria of a representative experiment are shown. The values for the induction of the zinc reporter were derived from at least three independent experiments. Host cells were infected at MOI 5 with STM strains as indicated, each containing the zinc reporter p5003. HeLa cells (C-E) and RAW macrophages (F-H) were analyzed as described in **Figure 3**. Mean values and standard deviations of triplicates of a representative experiment from three biological replicates are shown. Statistical analyses are indicated as for **Figure 3**. E) HeLa cells were infected with STM WT harboring p5194 [P_*uhpT*_::DsRed P_*znuA*_::sfGFP] (blue) and p5003 [P_*EM7*_::DsRed P_*znuA*_::sfGFP] (red).

In RAW264.7 macrophages, induction of 30% and 20% of STM WT at 8 h and 16 h p.i., indicated Zn^2+^ availability of less than 1 μM (**Figure 4**F, black frame, G). A stronger deficiency for Zn^2+^ ions was observed at 16 h p.i. for STM Δ*ssaV* (about 30% induced), and for STM WT isolated from activated RAW264.7 macrophages (**Figure 4**H). All other strains showed lower sfGFP intensity, thus low induction of the reporter. We concluded an environment poor in zinc in RAW264.7 macrophages, but predominantly with Zn^2+^ levels higher than 1 μM.

### Biotin synthesis is upregulated by intracellular STM

STM has been reported to rely on biotin biosynthesis for intracellular replication (Shi et al., 2009), suggesting that biotin is limited within the SCV. Culture in PCN medium with less than 200 μM D-biotin led to induction of P_*bioA*_::sfGFP, resulting in X-mean sfGFP intensity above 450 RFI (**Figure 5**AB). For STM released from HeLa cells, about 70% and 90% of STM WT at 8 h and 16 h p.i., respectively, exhibited such sfGFP intensity or even higher levels. The population of intracellular STM Δ*ssaV* or Δ*sifA* were similar to STM WT (**Figure 5**C, black frame, D). Therefore, almost all STM experience biotin concentration lower than 200 μM during intracellular presence. Bacteria with access to cytosolic components, i.e. the P_*uhpT*_-positive subpopulation, exhibited only slightly lower induction of P_*bioA*_::sfGFP (**Figure 5**E indicated by yellow frame). STM released from RAW264.7 macrophages showed similar results, again ca. 90% of the population of STM WT and Δ*ssaV* showed an induction corresponding to biotin concentrations lower than 200 μM (**Figure 5**F, black frame, G). The activation of RAW264.7 macrophages with IFN-γ resulted in mixed populations for STM WT and Δ*ssaV* (**Figure 5**F, black frame). Microscopy of infected HeLa cells (**Figure 5**I) and RAW264.6 macrophages (**Figure 5**J) showed the induction of the biotin biosynthesis reporter, almost all bacteria showed P_*bioA*_::sfGFP induction but the sfGFP intensity of the bacteria very heterogeneously distributed (**Figure 5**I).

**Figure 5:**
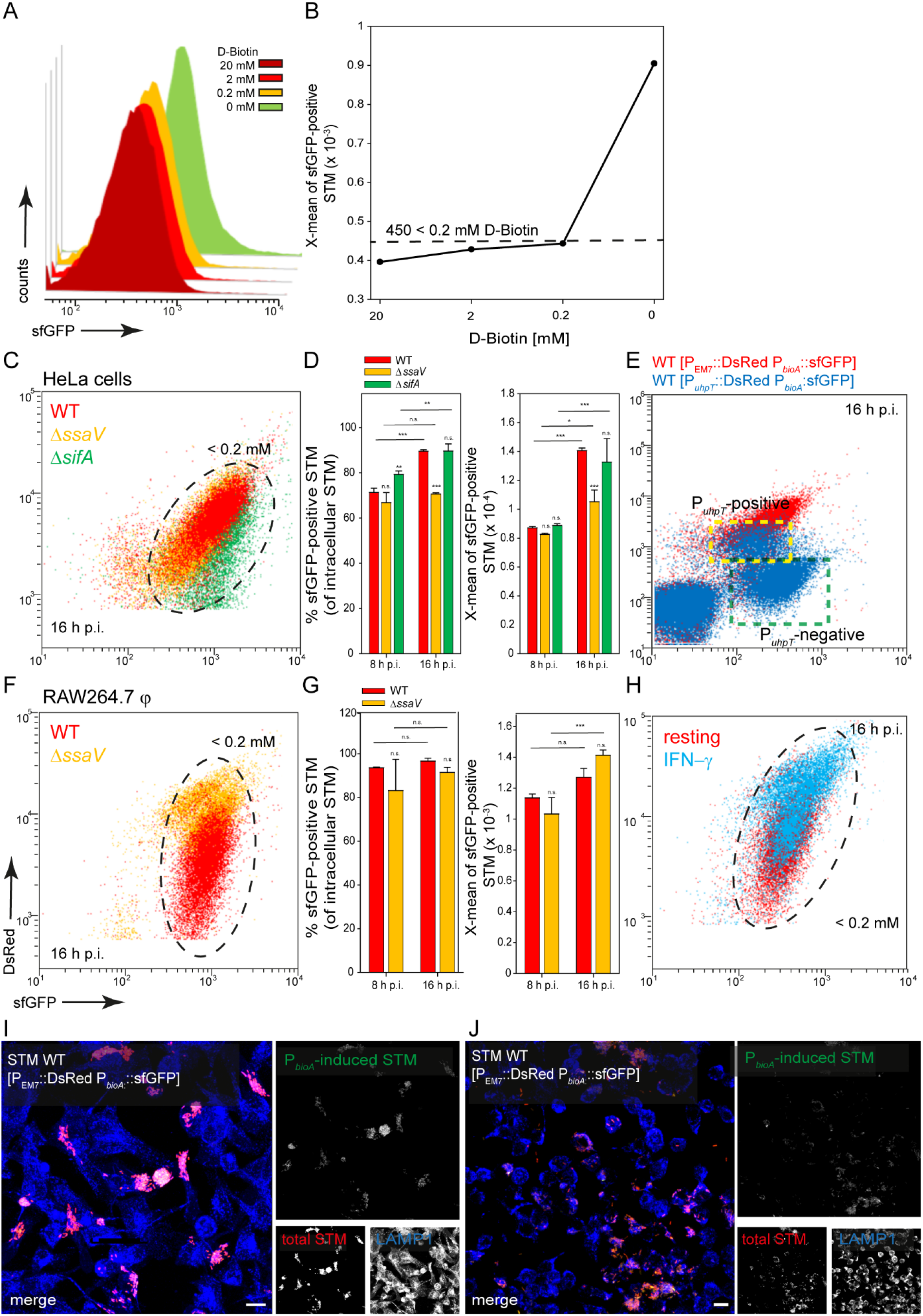
A dual fluorescence reporter for measuring the intracellular biotin synthesis of STM in HeLa cells or RAW264.7 macrophages. STM harboring p5067 for constitutive expression of DsRed, and sfGFP under control of P_*bioA*_ was cultured in in PCN minimal medium with various amounts of D-biotin. STM WT [p5067] was grown o/n in PCN (25), pH 7.4, diluted 1:31 in fresh PCN (1), pH 7.4 with various concentrations of D-biotin as indicated, and subcultured for 3.5 h. Samples were collected after 3.5 h (A) and X-means of sfGFP intensity of P_*bioA*_-induced bacteria was determined by FC (B). sfGFP intensities above 450 RFI indicate D-biotin concentrations lower than 200 μM. sfGFP intensities of P_*bioA*_-positive bacteria of a representative experiment are shown. The values for induction of the biotin reporter were derived from at least three independent experiments. Host cells were infected at MOI 5 with STM strains as indicated, each containing the biotin reporter p5067. HeLa cells (C-E) and RAW macrophages (F-H) were analyzed as described in **Figure 3**. Mean values and standard deviations from triplicates of a representative experiment from three biological replicates are shown. Statistical analyses are indicated as for **Figure 3**. E) HeLa cells were infected with STM WT harboring p5191 [P_*uhpT*_::DsRed P_*bioA*_::sfGFP] (blue) and p5067 [P_EM7_::DsRed P_*bioA*_::sfGFP] (red). Imaging for HeLa cells (I) and RAW264.7 macrophages were performed as described for **Figure 2**. Scale bars, 10 μm.

Addition of biotin to the medium of infected host cells reduced induction of the biotin reporter. In HeLa cells as well as in RAW264.7 macrophages, externally supplied biotin was internalized and can be used by the intracellular bacteria (**Figure 6**). We conclude that biotin is limited in the SCV of HeLa cells and macrophages, and for cytosolic STM in HeLa cells. Biotin synthesis is an important factor for survival and proliferation of intracellular bacteria.

**Figure 6:**
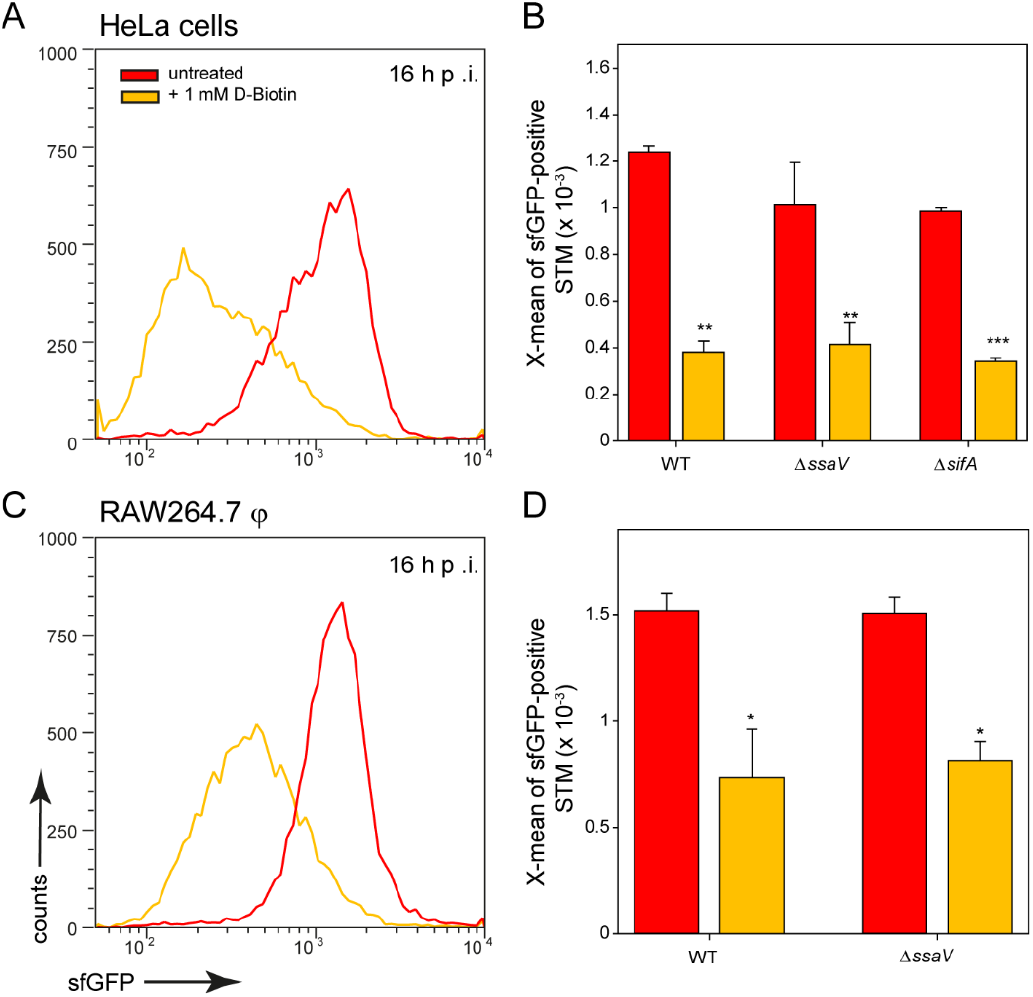
Changes in intracellular biotin availability for STM in HeLa cells or RAW264.7 macrophages. HeLa cells or RAW264.7 macrophages were infected at MOI 5 with STM WT, Δ*ssaV*or Δ*sifA* as indicated, each containing the biotin reporter p5067. 1 mM D-biotin (orange) was added 1 h p.i. to STM-infected HeLa cells or RAW264.7 macrophages, and maintained for the rest of the infection period. Host cells were lysed 16 h p.i., released STM were fixed, and subjected to FC to quantify sfGFP intensities of P_*bioA*_-positive STM. Data for STM WT [p5067] in HeLa cells (A) or RAW264.7 macrophages (C) of a representative experiment are shown. Representative quantification of X-mean sfGFP intensities of for STM WT, Δ*ssaV* and Δ*sifA* in HeLa cells (B) and RAW264.7 macrophages (D) at 16 h p.i. Mean values and standard deviations of P_*bioA*_-positive bacterial populations from triplicates of a representative experiment from three biological replicates are shown. Statistical analyses are indicated as for **Figure 3**.

### Iron is most limiting in the cytosol of epithelial cells and within macrophages

After invasion in host cells, STM encounter an iron-restricted environment. As a result, STM have evolved a variety of high-affinity iron acquisition systems to extract iron from this limiting environment *(Bjarnason et al., 2003)*. One system is an ABC transporter with a periplasmic binding protein-dependent transport system that is specific for metal ions, encoded by SPI1-located *sitABCD (Zhou et al., 1999; Zaharik et al., 2004; Ikeda et al., 2005)*. The promoter of *sitA* was used to investigate the iron concentrations available for STM in host cells. Because transcription of the *sit* operon dependents on oxygen content and growth phase *(Ikeda et al., 2005)*, the sfGFP intensity of the reporter was determined in growing LB cultures (**Figure 7**A). There was a strong induction of the reporter in an o/n culture, and the sfGFP signal dilutes in a growing subculture, but increases again in the late stage (6 h). After subculture for 2 h or 4 h, sfGFP intensity was lowest (**Figure 7**A). When iron chelator 2,2-dipyridyl was added to STM subcultured for 8 h, a particularly strong induction was observed, comparable to that of o/n cultures (**Figure 7**B). Transcription of *sitABCD* is negatively controlled by both MntR and Fur *(Ikeda et al., 2005)*, therefore induction of o/n cultures can be inhibited by addition of FeCl_2_ and MnCl_2_ (**Figure 7**CD). The pre-induction of STM after 8 h subculture was reduced by the addition of 100 μM FeCI_2_ or MnCI_2_ (**Figure 7**B). However, the lowest induction was achieved by combination of both metals. Addition of various concentrations of both ions led to concentration dependent decrease of P_*sitA*_::sfGFP induction. For STM exhibiting sfGFP intensities above 4,000 RFI, an intracellular environment with iron (Fe) and manganese (Mn) concentration below 100 nM can be deduced.

**Figure 7:**
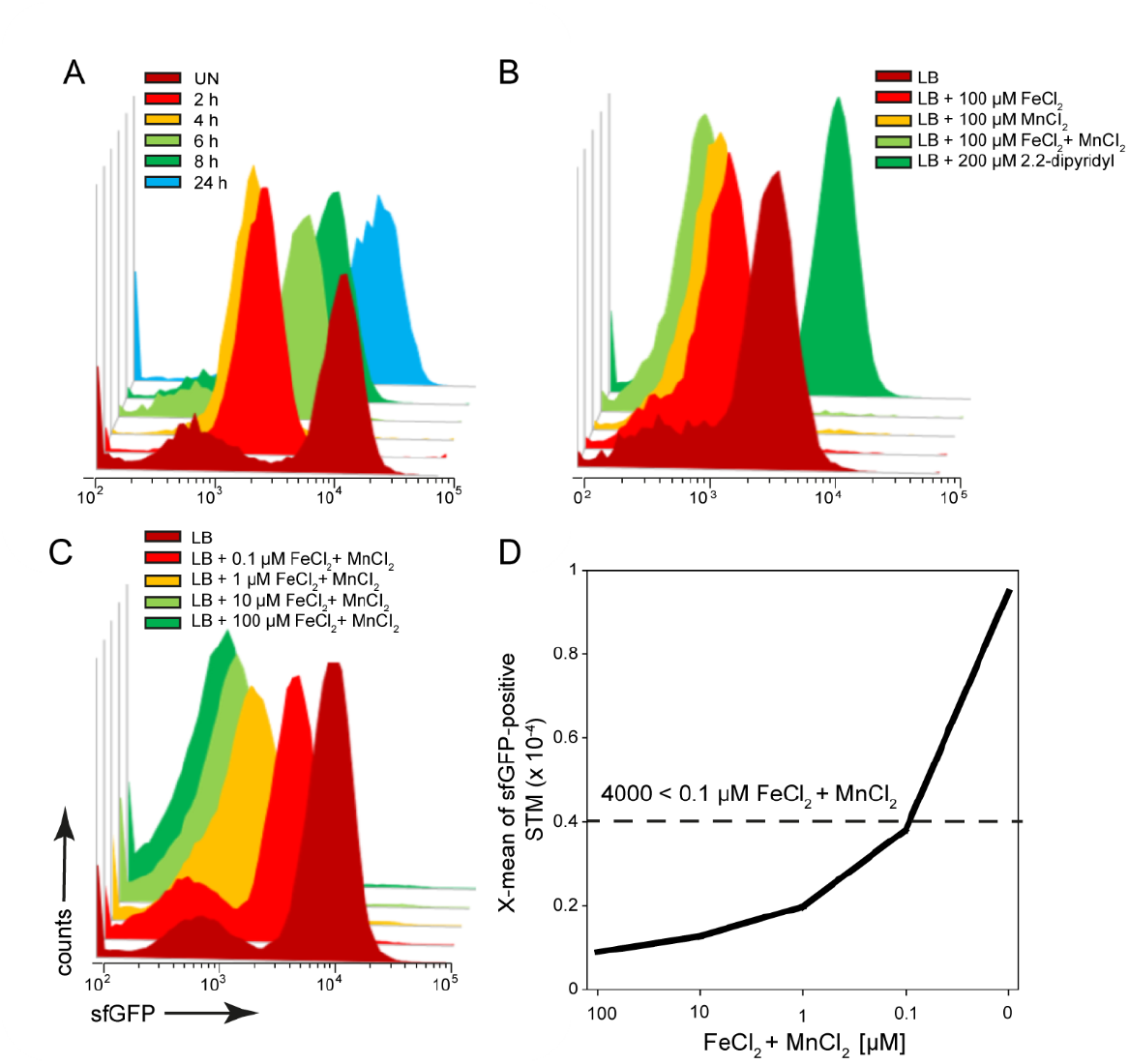
A dual fluorescence reporter for measuring iron limitation. STM harboring p5008 for constitutive expression of DsRed, and sfGFP under control of P_*sitA*_ was cultured in LB media. Induction of P_*sitA*_::sfGFP *in vitro* was determined by FC. A) STM WT [p5008] was grown o/n in LB, diluted 1:31 in fresh LB and subcultured. Samples were collected after various timepoints of subculture as indicated. B) WT [p5008] was grown o/n in LB, diluted 1:31 in fresh LB without or with addition of 100 μM FeCI_2_, and/or MnCI_2_, and 200 μM 2.2-dipyridyl, and subcultured for 8 h. C, D) STM WT [p5008] was grown o/n in LB, or LB supplemented by various concentrations of FeCI_2_ and MnCl_2_ as indicated. Samples were collected after 8 h of culture (C) and X-means of sfGFP intensity of P_*sitA*_-induced bacteria was determined by FC (D). sfGFP intensities above 4,000 RFI indicate FeCI_2_ + MnCI_2_ concentrations lower than 100 nM. sfGFP intensities of P_*sitA*_-positive bacteria of a representative experiment are shown. Concentrations for induction of the reporter were derived from at least three independent experiments.

STM WT, Δ*ssaV*, or Δ*sifA* strains harboring the P_*sitA*_ reporter were used to infect HeLa cells (**Figure 8**), or RAW264.7 macrophages (**Figure 9**). In HeLa cells, populations of STM with distinct levels of induction were observed. At 8 h p.i., ca. 90% of STM (subpopulation P2) showed strong induction comparable to *in vitro* growth at Fe^*2+*^ and Mn^*2+*^ concentrations less than 100 nM. At 16 h p.i., subpopulation P2 decreased (ca. 20% of bacteria < 100 nM) while P1 increased (ca. 80% of bacteria < 1 μM) for STM WT and Δ*ssaV*. Whereas concentrations below 100 nM are still available in the environment of STM and Δ*sifA* at 16 h p.i. (**Figure 8**AB, black frame, C). STM Δ*sifA* mutant has a predominantly cytosolic lifestyle, it can already be assumed that Fe^2+^ and Mn^2+^ ions are less present in the cytosol. Application of reporter P_*sitA*_::sfGFP in combination with P_*uhpT*_::DsRed for cytosolic presence clearly showed that the P_*uhpT*_-positive population (cytosolic) was stronger induced for P_*sitA*_ (**Figure 8**D, indicated by yellow frame). Imaging of infected cells also demonstrated the correlation between bacterial load of HeLa cells and iron limitation, since host cell with hyper-replication of STM showed high sfGFP intensities (**Figure 8**E).

**Figure 8:**
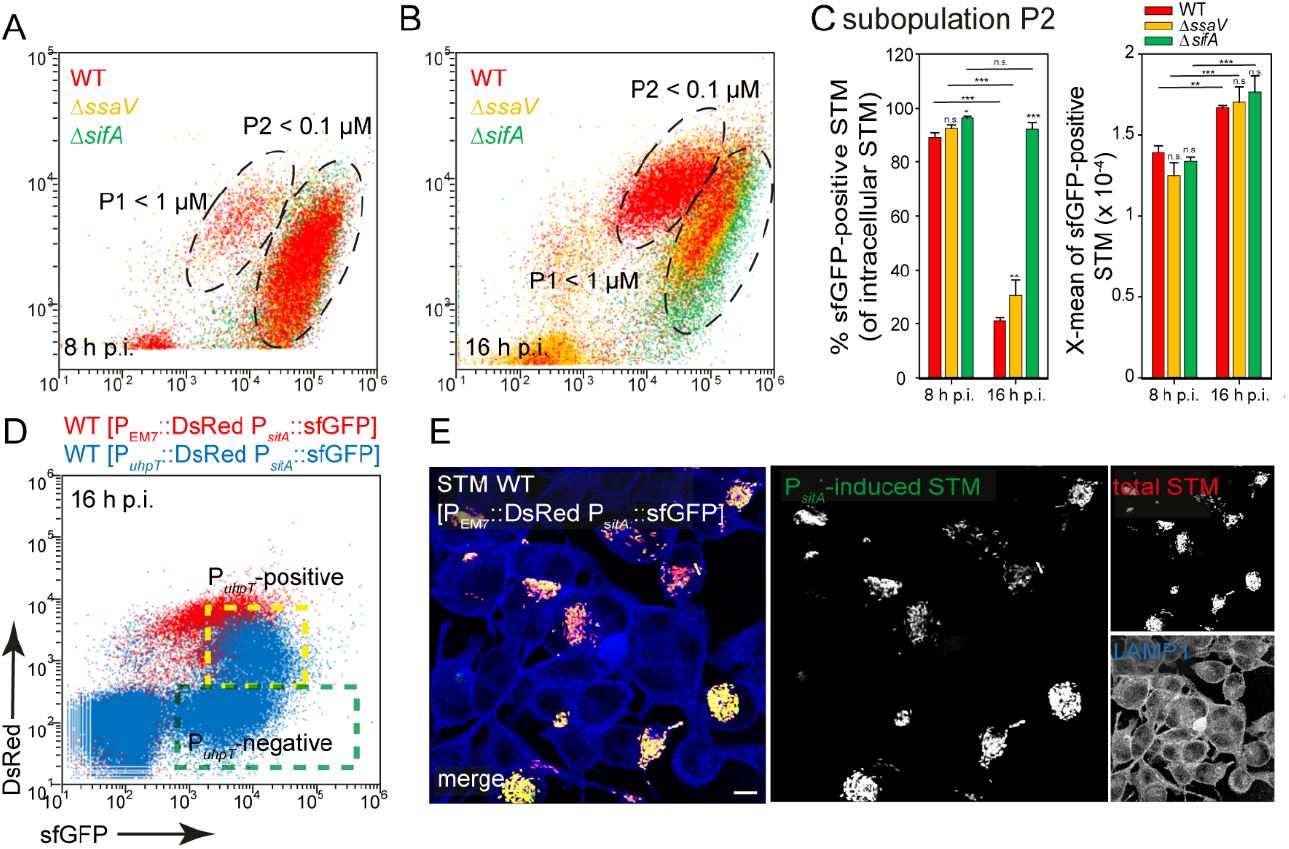
Determination of iron limitation of intracellular STM in HeLa cells. A, B) Infection of HeLa ells with STM strains as indicating, each containing the iron reporter p5008 was performed as described for **Figure 3**. Mean values and standard deviations from triplicates of a representative experiment from three biological replicates are shown. Statistical analyses are indicated as for **Figure 3**. E) HeLa cells were infected with STM WT harboring p5450 [P_*uhpT*_::DsRed P_*sitA*_::sfGFP] (blue), or p5008 [P_EM7_::DsRed P_*sitA*_::sfGFP] (red). Imaging was performed as described for **Figure 2**. Scale bar, 10 μm.

**Figure 9:**
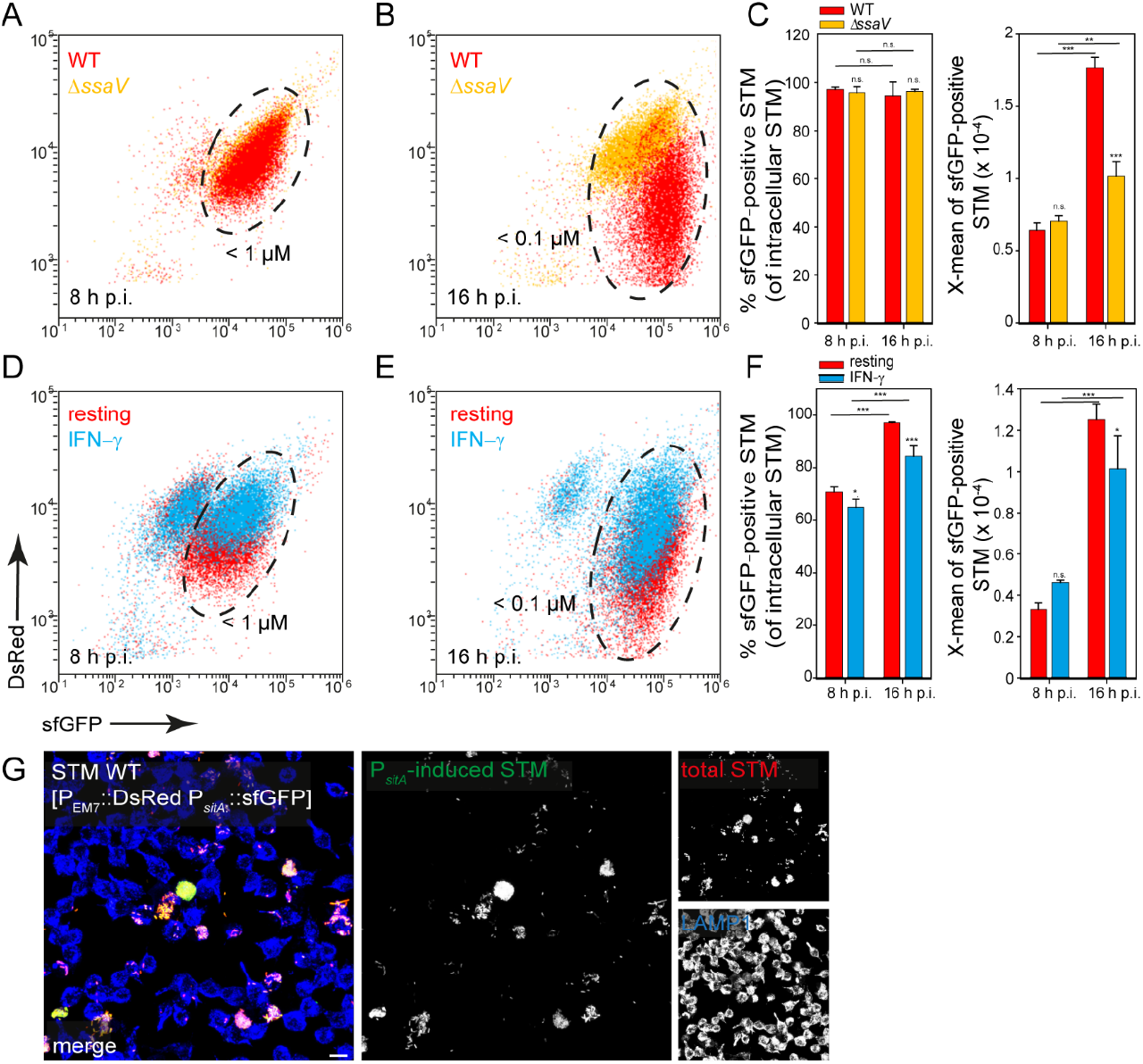
Determination of iron limitation of intracellular STM in RAW264.7 macrophages. Infection of RAW264.7 macrophages with STM strains as indicated, each containing the iron reporter p5008 was performed as described for **Figure 3**. Mean values and standard deviations from triplicates of a representative experiment from three biological replicates are shown. Statistical analyses are indicated as for **Figure 3**. Imaging was performed as described for **Figure 2**. Scale bar, 10 μm.

In contrast, bacteria isolated from RAW264.7 macrophages showed a strong induction of P_*sitA*_ at 8 h and 16 h p.i. (ca. 90% of the bacteria < 100 nM Fe^2+^/Mn^2+^ ions). The increase of sfGFP intensities from 8 h to 16 h p.i. indicated decreased Fe^2+^ and Mn^2+^ availability for intracellular STM over time of intracellular proliferation (**Figure 9**AB, black frame, C). Bacteria isolated from INF-γ-activated RAW264.7 macrophages revealed mixed populations for STM WT (**Figure 9**DE, black frame, F). The overview images clearly showed that a strong bacterial load led to a stronger induction of the iron reporter (**Figure 9**G). Thus, there is very little iron in the SCV, which decreased rapidly with bacterial proliferation.

An additional reporter was used to determine intracellular iron limitation. SufA is involved in the Fe-S cluster assembly and described to be induced in iron-limiting environments, and also by oxidative stress (Lee et al., 2004). The pre-induction of P_*sufA*_ in late cultures could not be stopped by addition of FeCI_2_ and/or MnCI_2_, only a 2-4 h subculture showed no pre-induction **Figure S 1**A). Nevertheless, P_*sufA*_-positive bacteria showed identical induced subpopulations like we found by P_*sitA*_-positive bacteria isolated within HeLa cells and RAW264.7 macrophages (**Figure S 1**BC). In addition, this reporter was induced stronger within IFN-γ activated RAW264.7 macrophages, possibly due to the higher exposure to oxidative stress (**Figure S 1**D).

In addition, a change in the culture medium of the host cells led to an altered reporter induction of STM (**Figure 10**). When iron was added to the medium of HeLa cells or RAW264.7 macrophages, STM showed a lower induction of the iron reporter. If iron was complexed by 2.2-dipyidyl, iron availability for STM also decreased, leading to a higher sfGFP intensity. In this way, STM adapts gene expression to the availability of iron and is able to successfully utilize the iron of the host cells and recruit it to SCV.

**Figure 10:**
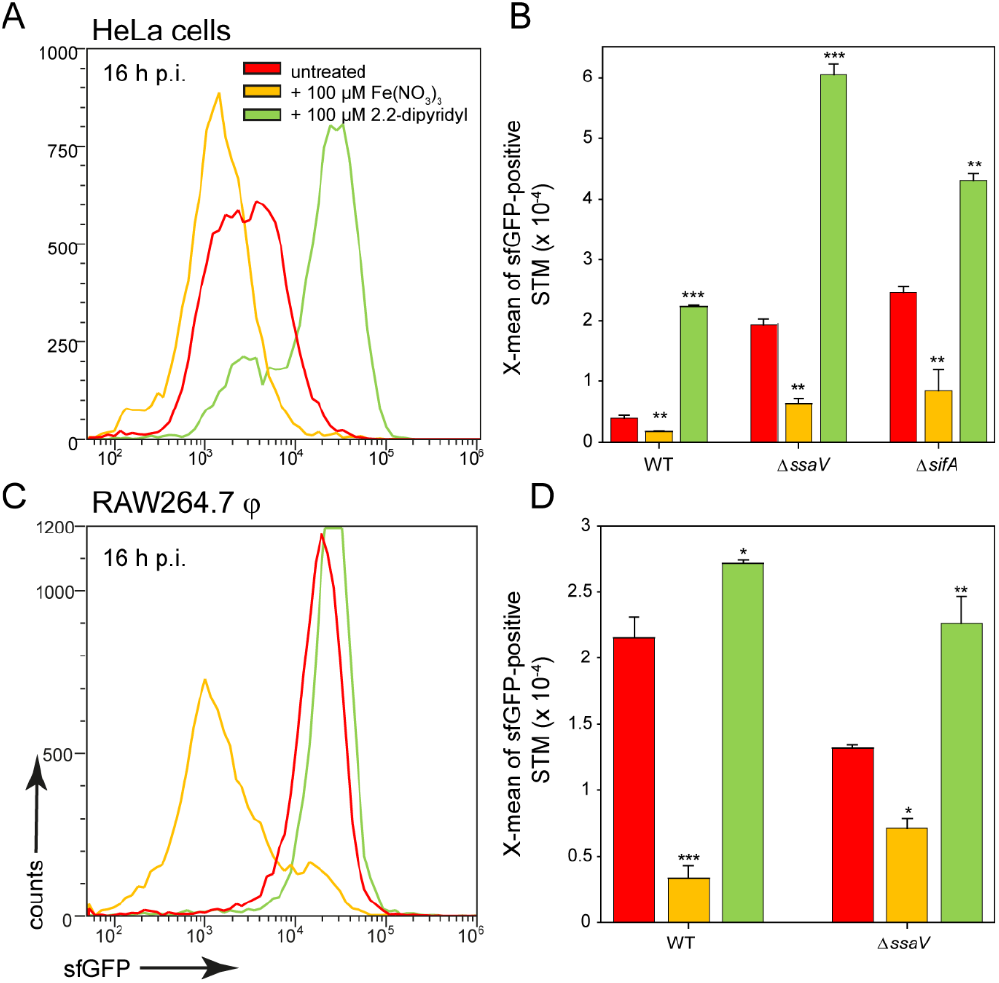
Effects of changes in iron availability in HeLa cells or RAW264.7 macrophages on intracellular STM. HeLa cells (A, B) or RAW264.7 macrophages (C, D) were infected at MOI 5 with STM WT, Δ*ssaV*, or Δ*sifA* strains as indicated, each containing the iron reporter p5008. At 1 h p.i., Fe(NO_3_)_3_ (orange) or 2,2-dipyridyl (green) were added to final concentrations of 100 μM to infected HeLa cells (A, B) or RAW264.7 macrophages (C, D) and maintained for the rest of the infection period. Host cells were lysed 16 h p.i., released STM were fixed, and subjected to FC to quantify sfGFP intensities of P_*sitA*_-positive STM. Data for STM WT [p5008] in HeLa cells (A) or RAW264.7 macrophages (C) of a representative experiment are shown. Representative quantification of X-mean sfGFP intensities for STM WT, Δ*ssaV*, or Δ*sifA* in HeLa cells (B) or RAW264.7 macrophages (D) at 16 h p.i. Mean values and standard deviations of P_*sitA*_-positive bacterial populations from triplicates of a representative experiment from three biological replicates are shown. Statistical analyses are indicated as for **Figure 3**.

### Correlation between bacterial burden and intracellular nutrient limitation

In some cases, a stronger nutrient limitation of intracellular STM WT compared to STM Δ*ssaV* was observed. The SCV-SIF continuum was suggested to increase access to endocytosed material from host cells, while STM in SCV without SIFs, such as the STM Δ*ssaV*, have reduced access to nutrient. STM within the SCV-SIF continuum also exhibit higher metabolic activity and replication **(Liss et al., 2017)**. To analyze how increased metabolic activity and replication affects the induction of nutrient reporters, we measured whole host cells infected with STM WT (**Figure 11**). FC was used to differentiate between infected (**Figure 11**, black frame) and non-infected host cells (**Figure 11**, grey frame), based on the constitutive DsRed signal of STM. In addition, a higher DsRed intensity also indicated a higher bacterial load of the host cells. The correlation of the DsRed signal with the sfGFP signal showed that sfGFP intensity increased with increasing bacterial load of the host cells (**Figure 11**, green frame). This means that there is a larger number of sfGFP-positive bacteria and/or bacteria with stronger sfGFP intensity in host cells. Reporter induction therefore also corelated to bacterial proliferation and metabolic activity. Comparing the DsRed intensity and the sfGFP intensity of whole cells between infected HeLa cells (**Figure 11**AB) and RAW264.7 macrophages (**Figure 11**CD) revealed DsRed signals much stronger in HeLa cells, indicating a generally higher bacterial load. In contrast, sfGFP intensities were higher in RAW264.7 macrophages, indicating generally higher nutrient restrictions.

**Figure 11:**
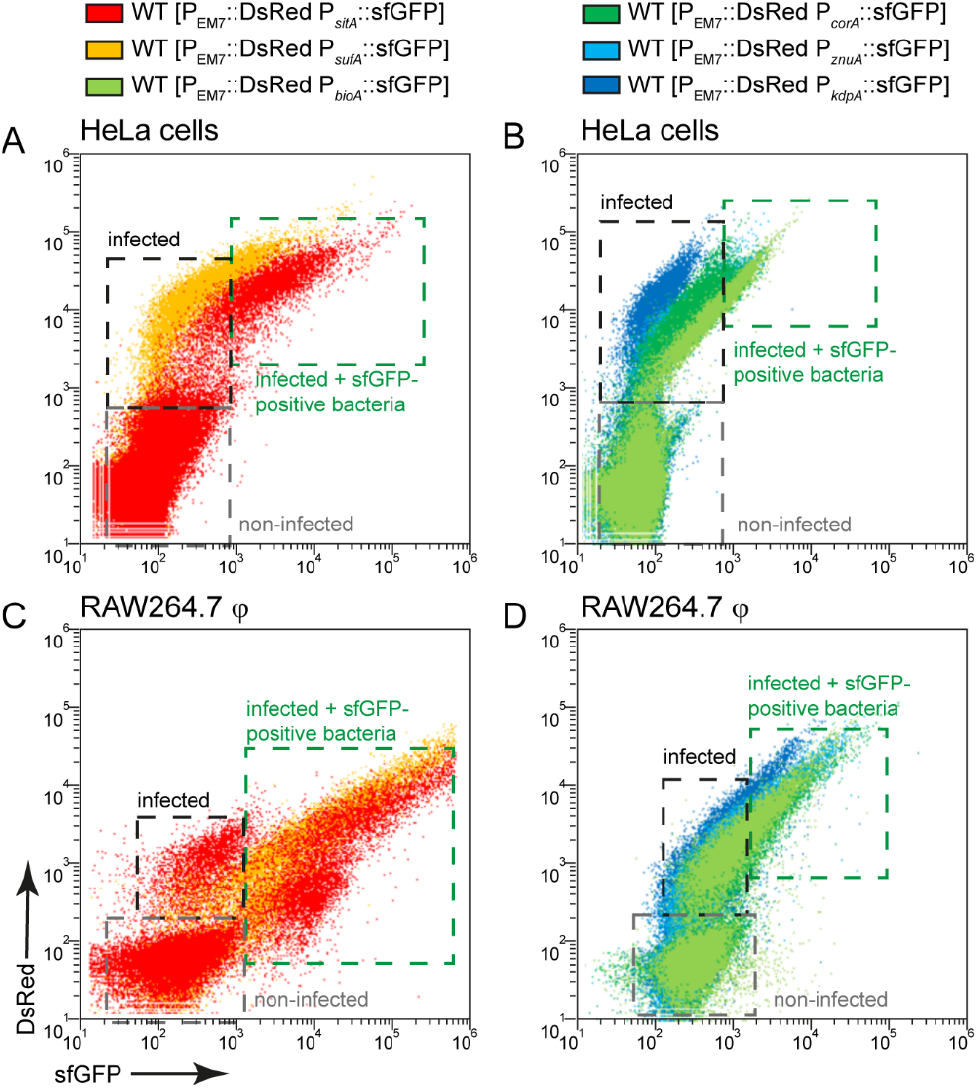
Correlation between bacterial burden and induction of various nutritional reporters. HeLa cells (A, B) or RAW264.7 macrophages (C, D) were infected with STM WT harboring p5008 [P_EM7_::DsRed P_*sitA*_::sfGFP], p5069 [P_EM7_::DsRed P_*sufA*_:sfGFP], p5067 [P_EM7_::DsRed P_*bioA*_::sfGFP], p5078 [P_EM7_::DsRed P_*corA*_::sfGFP], p5003 [P_EM7_::DsRed P_*znuA*_::sfGFP], or p5080 [P_EM7_::DsRed P_*kdpA*_::sfGFP] as indicated at MOI 5. The cells were detached and fixed 16 h p.i. Subsequently, the host cells were analyzed by FC to distinguish non-infected (grey frame), infected (black frame), and infected host cells with sfGFP-positive bacteria (green frame). Data for STM WT in HeLa cells (A, B) or RAW264.7 macrophages (C, D) of a representative experiment from three biological experiments are shown.

### Induction of nutritional reporters differs in macrophages isolated from human peripheral blood

Finally, induction of reporters was analyzed in human macrophages isolated from human Buffy Coat. It is known that STM cannot proliferate well in human macrophages compared to macrophages from STM-permissive mouse strains, such as the RAW264.7 cell line **(Lathrop et al., 2018)**. Nevertheless, STM is capable of causing systemic infection in humans. To better characterize the habitat of STM in primary human macrophages, the response of reporters was analyzed. In general, we observed that up to 40% of the bacterial population still showed detectable induction of sfGFP (**Figure 12**). The iron and biotin reporters showed the highest induction with about 40% induced intracellular bacteria, but sfGFP intensity was lower compared to bacteria isolated from RAW264.7 macrophages (**Figure 12**FG). The potassium reporter, like in RAW264.7 macrophages, was very poorly induced (**Figure 12**A). The magnesium reporter showed only a very low induction (**Figure 12**B), while the zinc reporter behaved similarly compared to induction in RAW264.7 (**Figure 12**C). An analysis of the bacterial load at 1 h and 8 h p.i. clearly showed that neither frequency of infected cells, nor the mCherry intensity human macrophages infected with STM [mCherry] increased (**Figure S 4**A). Within human macrophages, no bacterial replication, but rather bacterial survival was observed between 1-8 h p.i. The SPI2 reporter also showed a maximum induction in about 40% of the intracellular bacteria, but only 8.5% of the bacteria responded by measuring the metabolic activity using the Tet-inducible reporter at 8 h p.i. We conclude that 40% of the bacteria survived, probably mainly being in a persistent state with very low metabolic activity and low nutrient requirements. Limitations of Fe^**2+**^, Mn^**2+**^, Zn^**2**+^ ions and biotin were observed. However, there was no induction of the magnesium reporter in human macrophages, while potassium is present in sufficient quantities.

**Figure 12:**
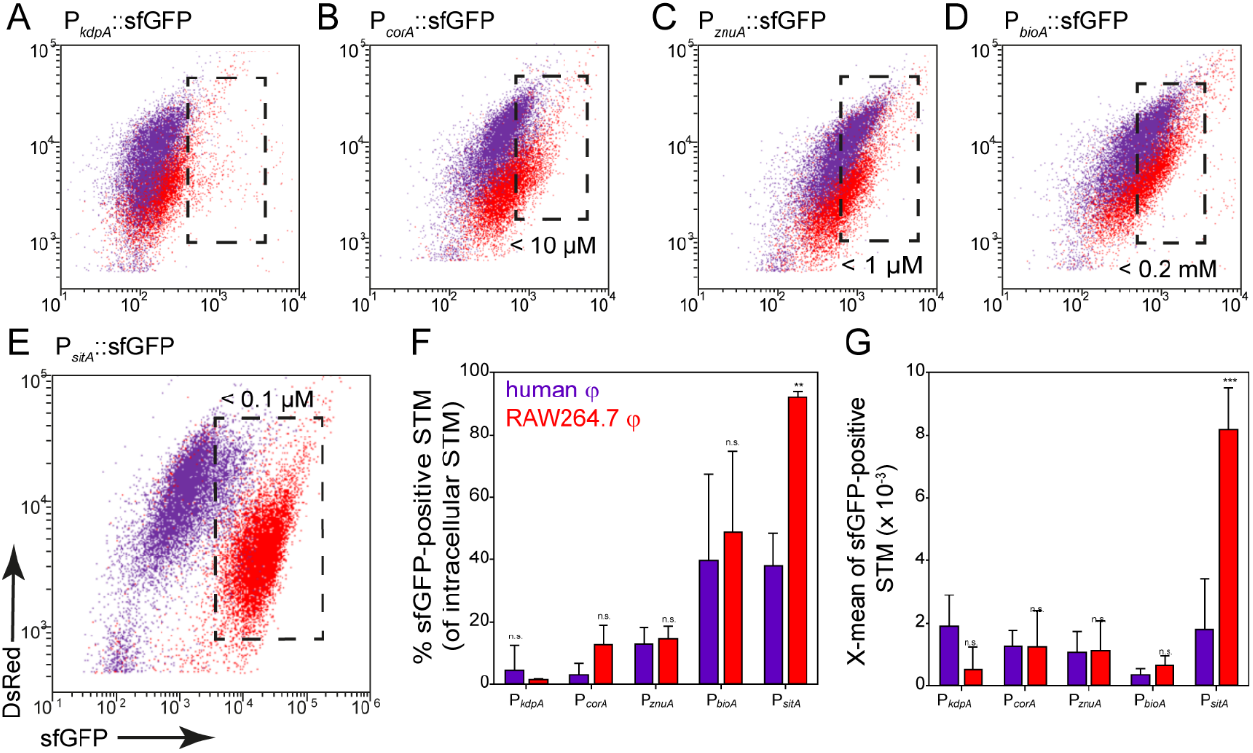
Nutritional limitations of STM in primary human macrophages. Human macrophages isolated from buffy coat (violet), or RAW264.7 macrophages (red) were infected at MOI 25 with STM WT strains harboring nutrient reporters as indicated. Human macrophages or RAW264.7 macrophages were lysed at 8 h p.i., released STM were fixed, and subjected to FC to quantify sfGFP intensities of sfGFP-positive STM (A, B, C, D, E). Representative quantification of population size and X-mean sfGFP intensities of sfGFP-positive population for STM WT in human macrophages (violet) or RAW264.7 macrophages (red) at 8 h p.i. (F, G). Shown are mean values and standard deviations of three biological replicates. Statistical analyses are indicated as for **Figure 3**.

## Discussion

We developed and applied an approach to study nutritional conditions in the SCV of host cells at single cell level. The use of dual fluorescence reporters allows conclusions about the intracellular concentrations of important nutrients. The data show that the nutrient composition varies in the habitats of different host cells, and reveals the adaptation of intracellular STM to nutritional limitations (summarized in **Figure 13**AB). Many studies have shown that STM can escape from the SCV and replicate efficiently in cytosol of epithelial cells, indicating a bimodal lifestyle (Malik-Kale et al., 2012). In macrophages, cytosolic bacteria are unable to survive due to clearance by xenophagy, or induction of pyroptosis **(Birmingham et al., 2006; Fink and Cookson, 2006)**. Therefore, we used HeLa cells to differentiate between nutrients in SCV and cytosol, but macrophages only to analyze the nutritional conditions of the SCV. The cytosol was already described as nutrient-rich habitat, resulting in a hyper-replicating phenotype of STM similar to a mutant strain lacking *sifA* (Beuzon et al., 2000). We conclude that Mg^2+^ and Zn^2+^ ions, and biotin are present in higher concentration, but Fe^2+^ and Mn^2+^ ions are more restricted in the cytosol of epithelial cells. By comparing reporter induction in the SCV of RAW264.7 macrophages and HeLa cells, we show that Fe^2+^ and Mn^2+^ ions are much more restricted in RAW264.7 macrophages. Biotin is present in similar concentrations, whereas K^+^ ions were limiting in a small bacterial subpopulation only in RAW264.7 macrophages. Mg^2+^ and Zn^2+^ ions show a heterogeneous availability, while these ions are more restricted at 8 h p.i. within RAW264.7 macrophages, concentration is more restricted within HeLa cells at 16 h p.i. For iron and biotin, we showed that bacteria are able to obtain these factors from the host cells in the SCV, as well as in cytosol. By adding iron or biotin to the external medium, induction of corresponding reporters decreased, indicating increased nutrient availability for intracellular STM. This finding is in line with the prior observations on access of STM to amino acids and glucose **(Popp et al., 2015; Liss et al., 2017)**. Furthermore, bacterial replication and thus bacterial burden of host cells influences the availability of nutrients. This is explained by the higher demand for nutrients by replicating bacteria, resulting in an interconnection between availability and consumption of nutrients, and replication of STM. Liss *et al*. (2017) already showed the influence of a large SCV-SIF continuum on STM access to endocytosed material. Together with our results, we conclude that bacteria in the cytosol have the best access to nutrients of the host cell, allowing fast proliferation. A large SIF-SCV continuum increases the frequency of fusion with vesicles, and allows the bacteria to replicate in the vacuolar compartment. In contrast, bacteria without SIF network are limited in nutrient recruitment, and remain unable to replicate (**Figure 13**C).

**Figure 13:**
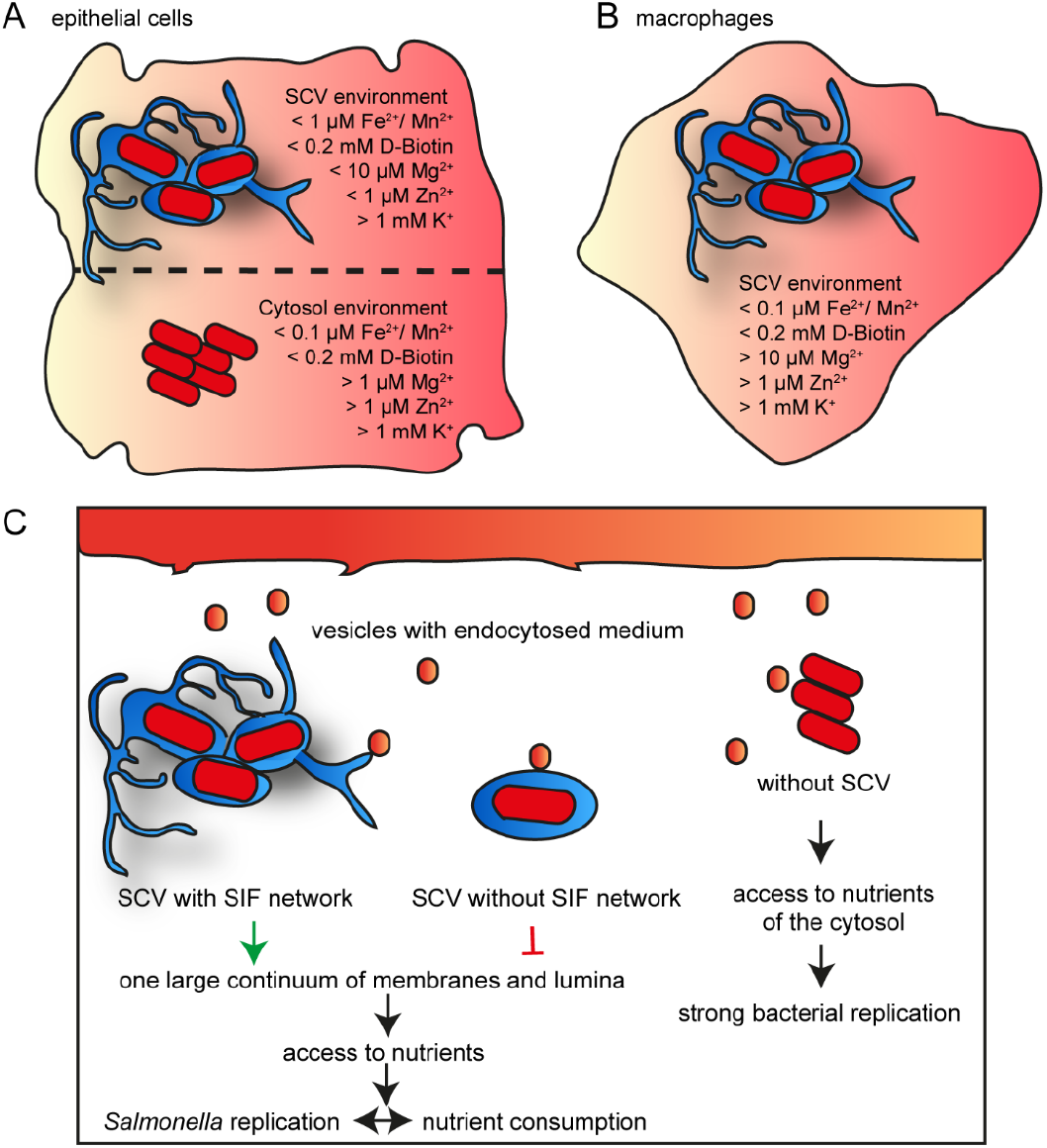
Model for nutrient availability of STM in distinct habitats of host cells. Comparison of availability of nutrients in epithelial cells between vacuolar and cytosolic STM (A), and within SCV in macrophages (B). Shown are values measured for the majority of the STM population at 16 h p.i. C) Nutrient access in various habitats of host cells.

Bacteria harboring fluorescent protein reporters are interesting tools to interrogate host cell environments on the level of single intracellular STM. To calibrate the reporters under *in vitro* conditions, we used batch cultures with various defined amounts of respective nutrients. The approach allows rapid determination of limiting concentration leading to induction. As cultural growth leads to continuous reduction of nutrient concentrations, the actual inducing concentration is likely lower for most nutritional factors. For a more detailed calibration, shift experiments with transfer in media with various amounts of nutrients be considered as performed for analyses of phosphate availability (Röder and Hensel, 2020b), or inhibition of bacterial growth. The presence of a second, non-regulated fluorescent protein has been used to identify in the intracellular bacterial population. However, constitutively expression of the slow maturing DsRed variant used here can also be deployed to measure rates of intracellular proliferation based on fluorescence dilution (Schulte et al., 2020).

Potassium was sufficiently available in the intracellular environment of STM and only a small subpopulation was induced within RAW264.7 macrophages. The *kdp* transport system was described to be induced by concentration lower than 5 mM K^+^ (Su et al., 2009), and we determined induction by less than 1 mM K^+^. This indicates high levels of potassium present in the SCV with concentration above 1-5 mM K^+^. Genes encoding transport systems for potassium were also not found to be induced in transcriptional profiles of intracellular STM in murine macrophages (Eriksson et al., 2003). Nevertheless, the three known K^+^ transporters were described to be critical for pathogenesis of STM in mice and chicks. Impaired K^+^ transport or deletion of transporters results in defect of the SPI1-T3SS (Liu et al., 2013). The low affinity K^+^ transporter Trk modulates the pathogenic properties of STM by increasing the expression and secretion of effector proteins of the SPI1-T3SS and enhancing epithelial cell invasion (Su et al., 2009). Potassium response and homeostasis also important for host colonization of *Mycobacterium tuberculosis* (MacGilvary et al., 2019). We conclude that K^+^ transport and maintaining K^+^ homoeostasis are important for extracellular and invasive strategies, but not for intracellular lifestyle of STM.

CorA is the primary Mg^2+^ transporter and STM lacking CorA showed attenuated virulence in murine or Caco-2 epithelial cell infection models (Papp-Wallace et al., 2008), but the mutant strain showed no growth defect and intracellular Mg^2+^ levels were not affected (Papp-Wallace and Maguire, 2008). Mg^2+^ is involved in virulence for example as extracellular Mg^2+^ concentrations mediate activation of the PhoP/PhoQ two-component system including the activation of SPI1, essential for invasion, and SPI2, essential for intracellular lifestyle (Garcia Vescovi et al., 1996; Deiwick and Hensel, 1999). Magnesium is the second most abundant cation and the concentration in mammalian cells range from 14-20 mM, while free concentrations are only between 0.5-0.7 mM (Romani, 2007). Within the SCV, Mg^2+^ concentrations of 10-50 μM were estimated by averaging the entire population (Garcia-del Portillo et al., 1992). Our single cell analyses indicated concentrations of less than 10 μM for 30% or 20 % for STM WT in HeLa cells or RAW264.7 macrophages, respectively, while 10-50 μM Mg^2+^ was available to the larger population of STM. Cytosolic bacteria showed lower sfGFP signals, indicating environments with 0.1-1 mM Mg^2+^. As host resistance factor SLC11A1 (a.k.a. NRAMP1) restricts STM proliferation through Mg^2+^ deprivation (Cunrath and Bumann, 2019), uptake by various transporters especially *Salmonella*-specific MgtB, appears to be important for intracellular Mg^*2+*^ homeostasis (Yeom et al., 2020).

ZnuA accumulates in low-zinc environments and is highly expressed by intracellular STM. Strains deficient in *znuA* are impaired in growth in Caco-2 epithelial cells and phagocytes (Ammendola et al., 2007). In mammalian cells, zinc is found as protein-bound or free ion. Concentrations of free zinc are in a picomolar range (Maret, 2017), suggesting that STM deploy zinc transporter ZnuABC to maximize zinc availability. We showed that strong *znuA* induction occurs at concentrations between 0-1 μM ZnSO^4^. We found that 50% of STM within the SCV in HeLa cells showed this level of induction, while higher concentrations are deduced for STM in host cell cytosol. Within RAW24.7 macrophages, the induction of STM WT decreased from 30% to 20% at 8 h and 16 h p.i., respectively. Removal of zinc is a host defense strategy against STM infection using by macrophages (Wu et al., 2017). STM-infected macrophages contained higher levels of free zinc than non-infected cells and cells that successfully eliminated the bacteria, suggesting that there is an bacteria-driven increase in intracellular zinc levels which weakens the antimicrobial defense and the ability of macrophages to eliminate the pathogen (Wu et al., 2017). Therefore, limiting cytosolic zinc levels may help to control infection by intracellular bacteria, but STM is able to increase the intracellular zinc concentration by an unknown mechanism (Wu et al., 2017). However, in HeLa cells the availability of zinc continues to decrease, maybe as result of the strong proliferation of STM.

Biotin also appears to be an important factor for intracellular life of STM, as almost all bacteria showed induction of the biotin reporter. We deduce that less than 200 μM biotin is available in the intracellular environment. Western blot and proteomic results showed increase upregulation of biotin biosynthesis of STM within macrophages (Shi et al., 2009), and reduced replication of a mutant strain was reported (Denkel et al., 2013). Biotin is synthesized only by microorganism and plants and is taken up in mammalian cells in a Na^+^-dependent manner (Said, 2009). We also demonstrated that biotin is only synthesized by STM if biotin is limiting in the SCV or cytosol, otherwise it is retrieved from the host. Induction of *bioA* of *S. flexneri* (Runyen-Janecky and Payne, 2002) also suggests that free biotin is limiting in eukaryotic host cells, since in *E. coli* these genes are repressed by 120 nM biotin (Eisenberg, 1985). Biotin limitation also affects other pathogens. In Malaria parasites, host biotin is required for liver stage development, and in *Francisella tularensis* intracellular biotin limitation triggers rapid phagosomal escape (Napier et al., 2012; Dellibovi-Ragheb et al., 2018). Upregulation of genes for biotin biosynthesis is also known for *Mycobacteria tuberculosis* in macrophages (Sassetti and Rubin, 2003; Woong Park et al., 2011). Inhibition of biotin biosynthesis is effective against antibiotic-resistant pathogens (Carfrae et al., 2020), underlining the importance of biotin biosynthesis for intracellular pathogens.

Iron and manganese are essential elements for mammalian cells as well as for bacteria. Only less than 100 nM Fe^2+^ and Mn^2+^ were determined in the SCV of RAW264.7 macrophages, or in the cytosol of HeLa cells. Within the SCV of MDCK cells a prior study postulated free Fe^2+^ concentration of 1 μM (Garcia-del Portillo et al., 1992), comparable to the concentration we found in the SCV of HeLa cells. Transcription of *sitABCD* is negatively controlled by both MntR and Fur *(Ikeda et al., 2005)*, therefore, we cannot distinguish between Fe^2+^ or Mn^2+^ concentrations in the SCV or the cytosol. Induction of *sitA* inside the intracellular environment was already described for STM (Liu et al., 2015), but deletion of *sitA* did not attenuate virulence (Ikeda et al., 2005), likely due to redundancy of high-affinity iron and manganese acquisition systems in STM. Inside host cells, iron is bound to specific proteins such as transferrin, lactoferrin and ferritin, resulting in availability of relatively low amounts of free iron. Also, Mn^2+^ concentrations in humans are very low ranging from 10 nM to 10 μM in serum and liver, respectively (Keen et al., 2000; Rahil-Khazen et al., 2002).

Acquisition of Fe^2+^ and Mn^2+^ is critical for pathogens to survive in the host, and pathogens have evolved sophisticated strategies to acquire and utilize host Fe^2+^ and Mn^2+^ (Ratledge and Dover, 2000; Juttukonda and Skaar, 2015). Fe^2+^ and Mn^2+^ restriction also play an important role in immune responds of the host. SLC11A1 is a transporter in phagosomes that actively limits Fe^2+^ and Mn^2+^ availability for STM in the SCV (Nairz et al., 2009). To counteract, STM is optimal adapted to these limiting conditions by procession of various transporters. The limitation of Fe^2+^ and Mn^2+^ ions is a defense strategy of the host against pathogens, but also provision of Fe^2+^ and thus Fe-induced or Mn-induced toxicity was described for *Leishmania* or *Neisseria meningitidis*, respectively (Veyrier et al., 2011; Silva-Gomes et al., 2013). Microbial iron acquisition depends on multiple, complex steps, many of them are not shared by higher eukaryotes. These acquisition systems are potential targets for the development of new antimicrobial therapies, as no effect on the host is anticipated (Ballouche et al., 2009). Furthermore, iron plays an essential role in the regulation of virulence genes. The low concentration of iron present in the host is an important signal to induce the expression of a wide variety of bacterial toxins and other virulence determinants in many different pathogens, such as *Shigella, Yersinia, Vibrio*, or *Neisseria* species and many more (Litwin and Calderwood, 1993). In *Legionella pneumophila* iron limitation triggers the early release into the host cell cytosol (O’Connor et al., 2016). Thus, there is a strong interplay between metabolism of iron and virulence.

The understanding of adaptation of STM to human macrophages and STM is still limited, thus we investigated the availability of nutrients in human monocyte-derived primary macrophages. STM does not replicate efficient in primary human macrophages (Schwan et al., 2000). In more detail STM did not proliferate in M1 macrophages (classically activated), while proliferation was possible in M2 (alternatively activated), or M0 (non-activated) macrophages (Lathrop et al., 2018). We used the growth factor GM-CSF for differentiation and predominantly obtained M1 macrophages. As induction of reporters for iron, biotin and zinc was observed in a small population of STM between 1-8 h p.i., we conclude limitation of these nutrients in the SCV of human M1 macrophages, and sensing and response of STM to these limitations.

In conclusion, STM shows a heterogenous adaptation to various host cells and habitats in regard to the nutrient acquisition. The formation of an extensive SIF network but also the expression of various genes for nutrient uptake or biosynthesis are important for the intracellular supply of nutrients. The use of fluorescent reporter is a suitable tool to characterize the vacuolar environment at single cell level to analyze heterogeneity and adaptation of intracellular pathogens.

## Acknowledgements

This work was supported by the Deutsche Forschungsgemeinschaft by grant HE 1964/18-2 and BMBF by grant 031L0093A as part of the Infect-ERA cluster SalHostTrop. We like to thank Monika Nietschke and Ursula Krehe for excellent technical support

## Authors contributions

JR and MH conceived the study, JR and PF performed experimental work, JR, PF, and MH analyzed the data, JR and MH wrote the manuscript.

## Materials and Methods

### Bacterial strains and growth conditions

*Salmonella enterica* serovar Typhimurium (STM) strains NCTC12023 (identical to ATCC 14028) and isogenic mutant strains are summarized in Table 1. STM strains were routinely cultured in Luria-Bertani (LB) broth containing 50 μg x ml^-1^ carbenicillin (Roth) if required for the selection of plasmids. Bacterial cultures were routinely grown in glass test tubes at 37 °C with aeration in a roller drum at ca. 60 rpm. For invasion of HeLa cells, fresh LB medium was inoculated 1:31 with overnight (o/n) cultures and incubated to 3.5 h with agitation in a roller drum. To test the induction of reporters *in vitro*, LB, PCN or MKM media were modified in various ways as indicated.

**Table 1.** Bacterial strains used in this study

| 1041 | <u>Designation</u> | <u>relevant characteristics</u> | <u>reference</u> |
| --- | --- | --- | --- |
| 1042 | NCTC 12023 | wild type | Lab collection |
| 1043 | MvP503 | NCTC 12023 Δ*sifA*::FRT | (Chakravortty et al., 2002) |
| 1044 | MvP1890 | NCTC 12023 Δ*ssaV*::FRT | noster (Noster et al., 2019) |

### Construction of plasmids

Plasmids used in this work are listed in Table 2. Oligonucleotides for generation of recombinant DNA molecules were obtained from IDT are specified in Table S1. The promoters of *sitA, sufA, bioA, corA, znuA*, or *kdpA* of STM were cloned as fragments of about 300 bp upstream of the translational start site. The fragments fused by Gibson assembly (GA) to sfGFP in plasmid p4889. This resulted in the dual fluorescence reporter harboring P_EM7_::DsRed for constitutive expression of DsRed and sfGFP under control of genes for acquisition of specific nutrients.

**Table 2.**
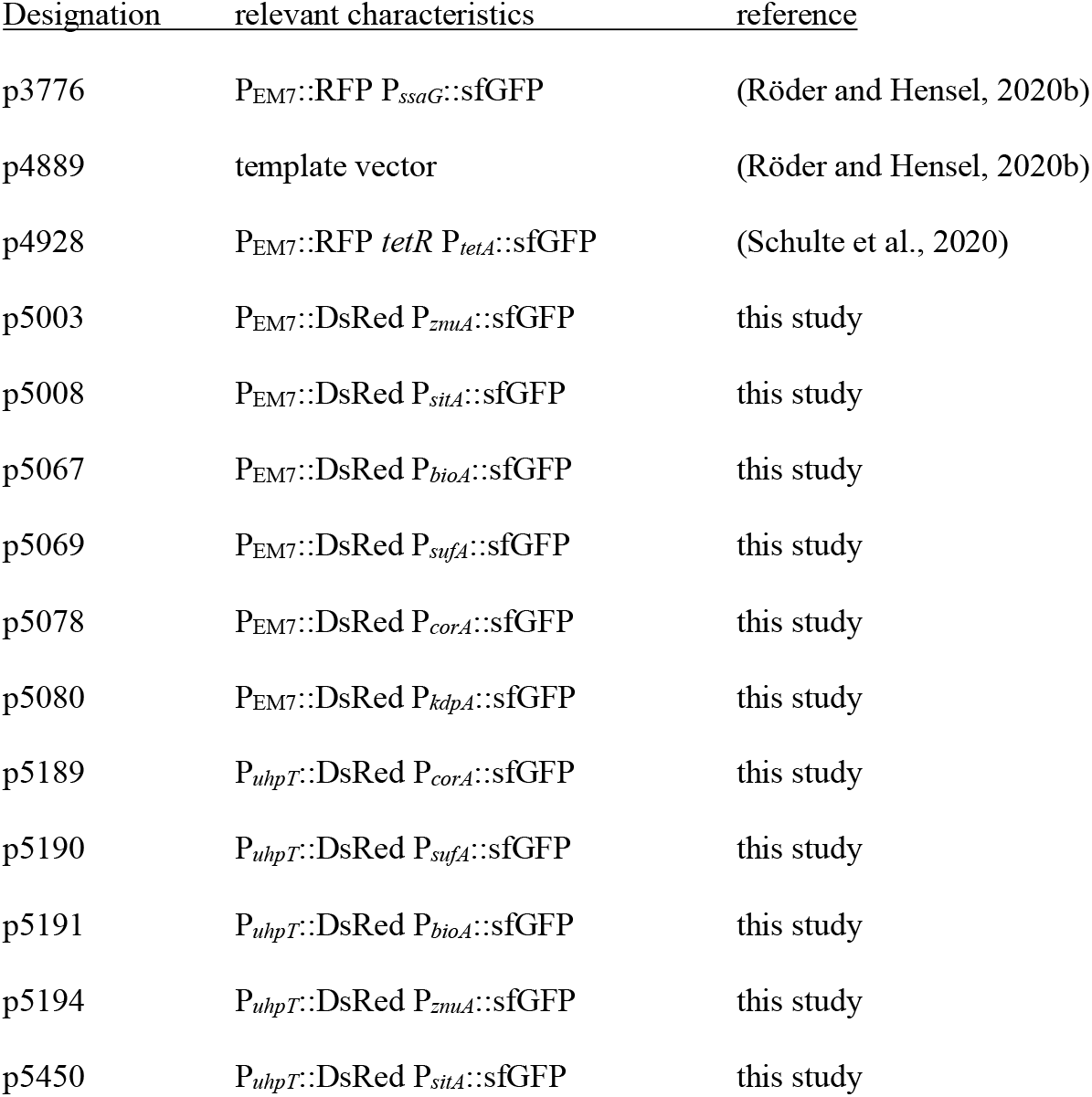
Plasmids used in this study

| 1047 | <u>Designation</u> | <u>relevant characteristics</u> | <u>reference</u> |
| --- | --- | --- | --- |
| 1048 | p3776 | P<sub>EM7</sub>::RFP P<sub>ssaG</sub>::sfGFP | (Röder and Hensel, 2020b) |
| 1049 | p4889 | template vector | (Röder and Hensel, 2020b) |
| 1050 | p4928 | P<sub>EM7</sub>::RFP *tetR* P<sub>tetA</sub>::sfGFP | (Schulte et al., 2020) |
| 1051 | p5003 | P<sub>EM7</sub>::DsRed P<sub>znuA</sub>::sfGFP | this study |
| 1052 | p5008 | P<sub>EM7</sub>::DsRed P<sub>sitA</sub>::sfGFP | this study |
| 1053 | p5067 | P<sub>EM7</sub>::DsRed P<sub>bioA</sub>::sfGFP | this study |
| 1054 | p5069 | P<sub>EM7</sub>::DsRed P<sub>sufA</sub>::sfGFP | this study |
| 1055 | p5078 | P<sub>EM7</sub>::DsRed P<sub>corA</sub>::sfGFP | this study |
| 1056 | p5080 | P<sub>EM7</sub>::DsRed P<sub>kdpA</sub>::sfGFP | this study |
| 1057 | p5189 | P<sub>uhpT</sub>::DsRed P<sub>corA</sub>::sfGFP | this study |
| 1058 | p5190 | P<sub>uhpT</sub>::DsRed P<sub>sufA</sub>::sfGFP | this study |
| 1059 | p5191 | P<sub>uhpT</sub>::DsRed P<sub>bioA</sub>::sfGFP | this study |
| 1060 | p5194 | P<sub>uhpT</sub>::DsRed P<sub>znuA</sub>::sfGFP | this study |
| 1061 | p5450 | P<sub>uhpT</sub>::DsRed P<sub>sitA</sub>::sfGFP | this study |

The promoter of P_*uhpT*_ of STM was cloned as 251 bp fragment upstream of the translational start site of *uhpT* and EM7 was replaced by P_*uhpT*_ using GA. The fragments fused by Gibson assembly (GA) to DsRed. This resulted in the double inducible reporter P_*uhpT*_::DsRed and sfGFP under control of genes for acquisition of specific nutrients.

### Cell lines and cultivation

Human epithelial cell line HeLa were maintained in DMEM containing 4.5 g x l^-1^ glucose, 4 mM L-glutamine and sodium pyruvate (Biochrom) supplemented with 10% FCS in an atmosphere of 5% CO_2_ and 90% humidity at 37 °C. The murine macrophage cell line RAW264.7 (ATCC no. TIB-71) were cultured in DMEM containing 4.5 g x 1^−1^ glucose and 4 mM stable glutamine (Biochrom) supplemented with 6% FCS.

### Preparation of primary human macrophages

Primary human macrophages were prepared from Buffy Coat from pooled blood samples of anonymous donors (obtained from the Deutsches Rotes Kreuz) as described in Bonifacino *et al*. (2004) Page 2/6 – 2/9 (Bonifacino et al., 2004). Preparation of lymphocytes by Ficoll-Hypaque gradient was performed as described, alternatively to whole blood, Buffy Coat was mixed 1: 1 with PBS. For differentiation into monocytes/macrophages, the isolated lymphocytes were thawed, seeded and maintained in RPMI-1640 (Biochrom), supplemented with 20% FCS and 2.5 ng x ml^-1^ GM-CSF (Peprotech). After 5-7 days, the purity of the monocyte/macrophage population was checked by staining with FITC anti-human CD14 antibody (BioLegend) and FC and used for infection.

### Infection of host cells

Host cell infections were performed as described previously (Rajashekar et al., 2008). Briefly, RAW246.7 macrophages and human macrophages were infected with o/n cultures and HeLa cells were infected with 3.5 h subcultures of STM with a multiplicity of infection (MOI) of 5 or 25. The bacteria were centrifuged onto the cells at 500 x g for 5 min, incubated for 25 min at 37 °C in an atmosphere of 5% CO_2_ before extracellular bacteria were removed by washing thrice with PBS. Subsequently, host cells were maintained in growth media containing 100 μg x ml^-1^ gentamicin for the rest of the experiment.

### Imaging

HeLa cells and RAW267.4 macrophages cultured in 24-well chamber slides on cover slips and infected at MOI of 5. The infection was performed as described above and 16 h after infection, Imaging by CLSM was carried out using a Leica SP5 system with environmental periphery.

### Quantitation by flow cytometry analyses

HeLa cells, RAW264.7 and primary human macrophages were infected at MOI of 5 or 25 for 25 min. 8 h and 16 h p.i., cells were lysed with 0.1 % Triton X-100 and released bacteria were fixed with 3% PFA for subsequent flow cytometry analyses using the Attune NxT Cytometer (Thermo Fischer). Alternative HeLa cells were detached with Biotase or RAW264.7 macrophages with ice-cold 5 mM EDTA in PBS. Infected and non-infected cells were fixed with 3% PFA for subsequent flow cytometry analyses using the Attune NxT Cytometer (Thermo Fischer).

Experiments were performed in triplicates at least three times. Data were analyzed with Attune NxT 2.5. Statistical analyses were performed using One Way Anova versus control with SigmaPlot 11 (Systat Software).

### Immunostaining

HeLa cells and RAW264.7 macrophages were infected with various STM strains as described, cells were fixed 16 h p.i. with 3% PFA for 10 min, washed with PBS and incubated in blocking solution (PBS containing 2% goat serum, 2% bovine serum albumin, 0.1% saponin freshly added from 3% stock solution in PBS). Permeabilized cells were stained with primary antibody mouse α hLAMP1, IgG1 monoclonal (DSHB) for 1 h, washed with PBS thrice, and detection was performed using secondary antibody goat α mouse Alexa 647, IgG (Fab) (Dianova) for 1 h.

## Supplementary Figure Legends

**Figure S 1:**
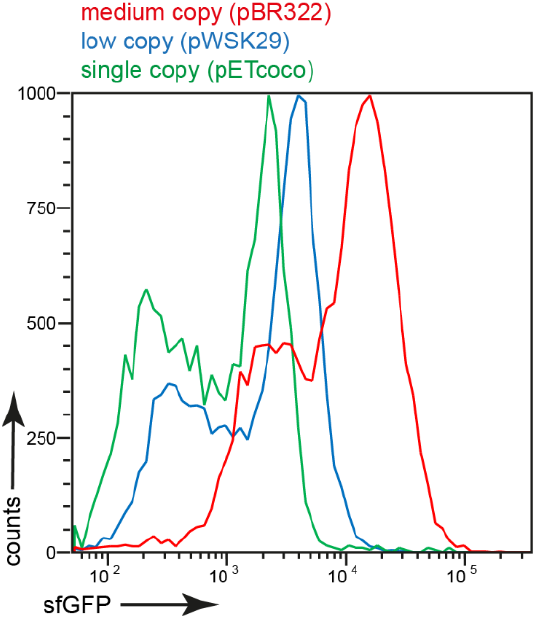
Comparison of induction of P_*sitA*_::sfGFP in plasmids with different copy numbers. HeLa cells were infected at MOI 5 with STM WT containing iron reporters p5008, p5235 or p5564 harboring P_*sirA*_::sfGFP in plasmid backbones with various copy numbers. The cells were lysed and fixed 16 h p.i. Subsequently, the bacteria were subjected to FC to quantify the sfGFP intensity of the P_*sitA*_-induced bacteria. Data from one biological replicate.

**Figure S 2:**
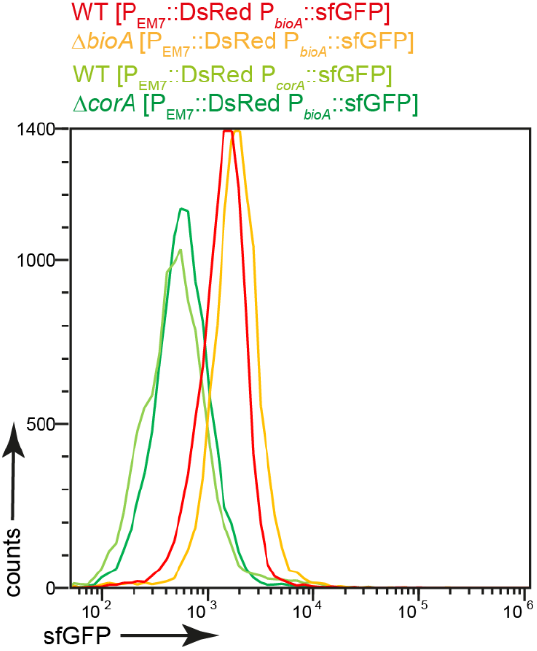
Comparison of reporter induction in STM WT and mutant strains. HeLa cells were infected at MOI 5 with STM WT and mutant strains deficient in *bioA* or *corA*, containing the reporter p5067 or p5078 as indicated. The cells were lysed 16 h p.i, STM released, and fixed. Subsequently, the bacteria were subjected to FC to quantify the sfGFP intensity. Data from one biological replicate.

**Figure S 3:**
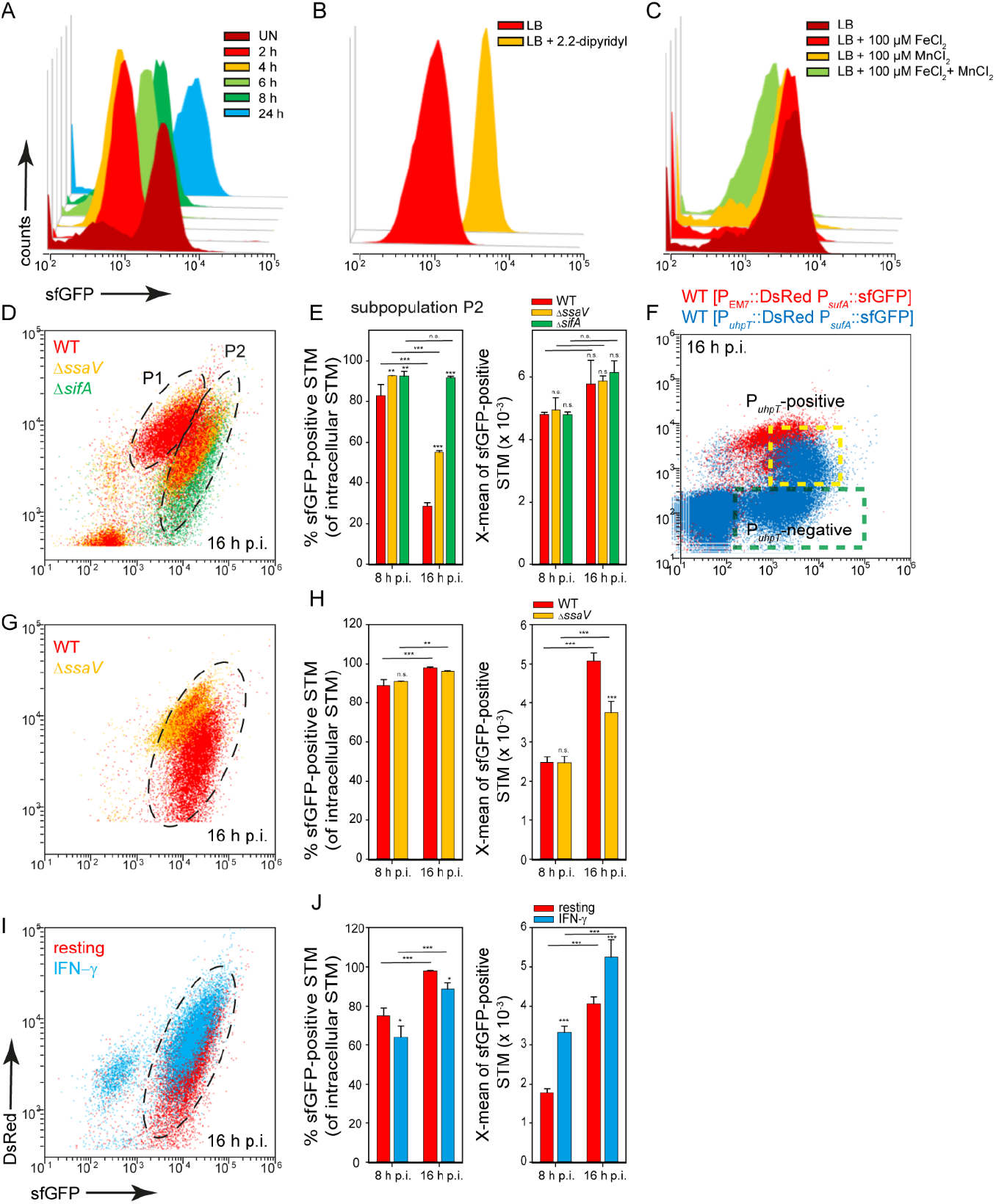
An alternative dual fluorescence reporter for measuring iron limitation. STM harboring p5069 for constitutive expression of DsRed, and sfGFP under control of P_*sufA*_ was cultured in LB media. Induction of P_*sufA*_::sfGFP *in vitro* was determined by FC. A) STM WT [p5069] was grown o/n in LB, diluted1:31 in fresh LB and subcultured for various time points as indicated. Samples were collected as indicated. B) WT [p5069] was grown o/n in LB, diluted 1:31 for 3.5 h in fresh LB without or with 200 μM 2.2-dipyridyl. C) WT [p5008] was grown o/n in LB, or LB complemented with FeCl_2_ and/or MnCl_2_ as indicated. Samples were collected after 3.5 h of culture. sfGFP intensities of P_*sufA*_-positive bacteria of a representative experiment are shown, the value for the induction of the reporter was derived from at least 3 independent experiments. D-J) Host cells were infected at MOI 5 with STM strains as indicated, each containing the iron reporter p5069. HeLa cells (D-F) and RAW macrophages (G-J) were analyzed as described for **Figure 3**. Mean values and standard deviations from triplicates of a representative experiment are shown. Statistical analyses are indicated as for **Figure 3**. E) HeLa cells were infected with STM WT harboring p5190 [P_*uhpT*_::DsRed P_*sufA*_:sfGFP] (blue) and p5069 [P_EM7_::DsRed P_*sufA*_:sfGFP] (red).

**Figure S 4:**
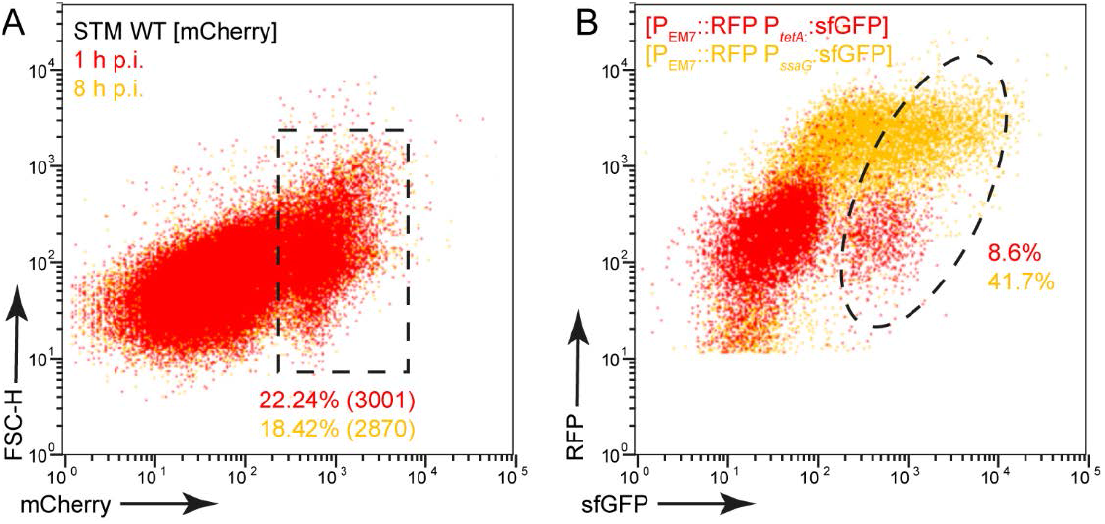
Intracellular proliferation, SPI2 gene expression, and metabolic activity of STM in human macrophages. A) Human macrophages were infected at MOI 5 with STM WT [mCherry]. The cells were detached and fixed at 1 h (red) or 8 h p.i. (orange). Subsequently, the macrophages were analyzed by FC. B) Human macrophages were infected at MOI 25 with STM WT [p4928] (red) or STM WT [p3776] (orange). For infection with STM [p4928], induction of the reporter was induced by addition of AHT to 100 ng x ml^-1^ at 6 h p.i., and cells were analysed 8 h p.i. STM was released, fixed and subsequently subjected to FC to quantify the sfGFP-positive bacteria. Data from one biological replicate.

## Suppl. Tables

**Table S1.**
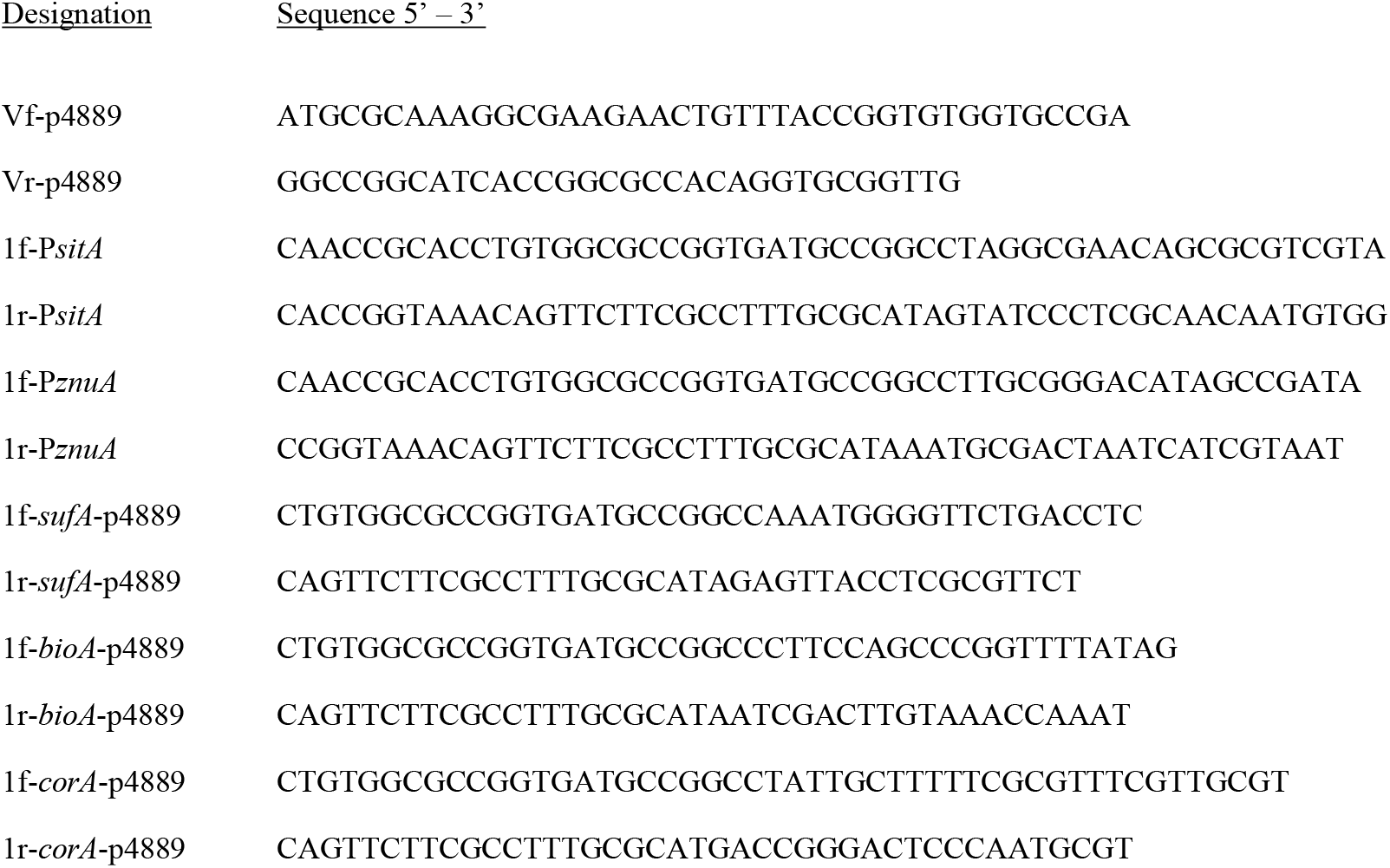
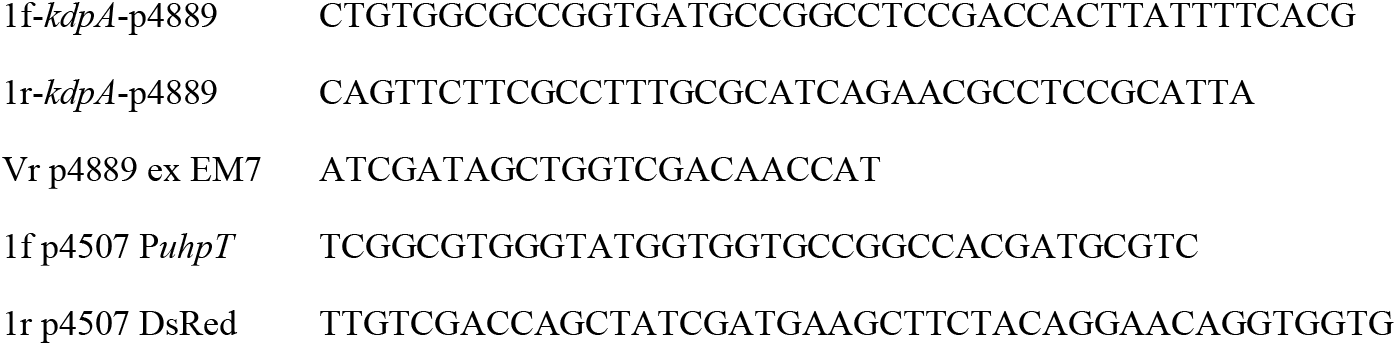
Oligonucleotides used in this study

